# Detecting and quantifying rare sex in natural populations

**DOI:** 10.1101/2025.06.03.657731

**Authors:** Tymoteusz Pieszko, Jerome Kelleher, Christopher G. Wilson, Timothy G. Barraclough

## Abstract

The distinction between sexual and asexual reproduction is fundamental to eukaryotic evolution. Testing theories about the evolution of reproductive modes first requires knowing whether sex is present or absent in a population. While this seems straightforward, the literature on asexuality reflects a history of shifting claims and uncertainty regarding reproductive mode, especially where sex is potentially rare or cryptic. Here, we develop a new framework to address the challenges in detecting and quantifying sexual reproduction from population genomic data. We first show that commonly calculated population genetic statistics do not reliably distinguish sexual and obligate asexual scenarios if asexuality is accompanied by sex-independent homologous recombination, as emerging evidence indicates is often the case. We then present a new method to quantify the relationship between evolutionary trees and mode of reproduction by classifying local trees for pairs of diploid individuals using ancestral recombination graphs (ARGs). This approach accurately discriminates signatures of genetic exchange and homologous recombination, though some uncertainty remains because of unavoidable biases in the steps needed to reconstruct trees from genome data. We introduce new statistics and simulation models to account for common reconstruction biases and demonstrate their utility by uncovering contrasting reproductive histories in the facultatively sexual budding yeast *Saccharomyces cerevisiae*. This approach offers the potential for improved quantitative inference of reproductive modes that can readily be extended to a broad range of eukaryotes.

**Significance Statement:** Determining how often, if at all, organisms have sex has implications across biology. Studies often use population genomic data to interrogate the private life of putative asexuals, but the results prove surprisingly inconclusive. We develop a framework to address the challenges in detecting and quantifying rates of sex. New simulation models show how sex-independent recombination, which occurs widely across a range of asexual eukaryotes, causes genetic patterns to resemble sexual populations even when sex is absent. An approach based on ancestral recombination graphs (ARGs) and classification of local trees accounts for these problems and quantifies the remaining uncertainty due to inevitable reconstruction biases. Our framework will enable improved inferences of reproductive mode across a wide range of eukaryotes.

## Introduction

Sex is a defining feature of eukaryotic life, and fundamental questions about its benefits, costs and prevalence remain of considerable interest in evolutionary biology (Hartfield & Keightley, 2012; Maynard Smith, 1978; Otto, 2021; Kondrashov, 2018). Testing hypotheses about the evolution of reproductive mode depends critically on the ability to detect the presence or absence of sex reliably and, if sexual and asexual reproduction co-occur, to estimate the frequency of sex. Because direct observations in the wild are often unfeasible (but see Neiman et al., 2018), a more common approach has been to use DNA sequence data to infer the occurrence and frequency of different modes of reproduction (Schurko et al., 2009; Tibayrenc et al., 1991), first from genetic markers (e.g., Burt et al., 1996; Kuhn et al., 2021) and now chiefly from whole genome data (Brandt et al., 2021; Freitas et al., 2023; Vakhrusheva et al., 2020; Van den Broeck et al., 2020).

Sexual reproduction has important consequences for genetic variation (Otto, 2003) and therefore patterns of sequence evolution were long predicted to provide clear evidence of reproductive modes (Maynard Smith, 1999; Normark et al., 2003; Schurko et al., 2009; Tibayrenc et al., 1991). In some taxa, this approach yielded insights that were later validated (Eitel et al., 2011; Signorovitch et al., 2005). However, numerous cases remain ambiguous, including some of agricultural (Lee et al., 2024; Yildirir et al., 2020) and medical (Carpenter et al., 2012; Cooper et al., 2007; Poxleitner et al., 2008; Bradic & Carlton, 2018) importance. The literature on asexual animals alone demonstrates the complexities in applying these methods, with numerous revised claims and uncertainty that spans decades of study (Boyer et al., 2021; Freitas et al., 2023; Mark Welch et al., 2008; Molinier et al., 2025; Schwander et al., 2011; Vakhrusheva et al., 2020). Classically predicted features of obligate asexuality (Normark et al. 2003) have failed to materialise in genomic surveys (Jaron et al., 2021; Nowell et al., 2018), while sequence-based evidence for the presence of cryptic sex sometimes contradicts the described natural history (Wilson et al., 2024). Before we can confidently assess reproductive modes, we need to understand why this remains such a difficult task.

One fundamental problem is that many tests do not formally separate the two main components of sexual reproduction – crossover recombination, and genetic exchange mediated by chromosome segregation and the syngamy of haploid gametes (Fig. 1A). It is now clear that homologous recombination between diploid chromosomes also occurs during asexual reproduction via a range of mechanisms, such as gene conversion, crossing-over, and break-induced replication associated with either mitotic or meiotic pathways (Jinks-Robertson & Petes, 2021; Engelstädter, 2017). We group such sex-independent mechanisms of recombination using the term ‘asyngamous recombination’ (Wilson et al., 2024). The main evidence for asyngamous recombination comes from the loss of heterozygosity (LOH) (Fig. 1B and C), observed even in lab-reared populations where the contribution of sex can be excluded. LOH is widespread across a range of asexual eukaryotes, including plants such as pineapple (Chen et al., 2019), fungi such as *Saccharomyces cerevisiae* (Sui et al., 2020), *Candida albicans* (Forche et al., 2011) and *Aspergillus nidulans* (Cardoso et al., 2010), diatoms (Bulankova et al., 2021), and parthenogenetic animals (Flynn et al., 2017; Houtain et al., 2024; Simion et al., 2021). Some forms of asyngamous recombination additionally result in reciprocal exchanges between homologous chromosomes (Fig. 1D) (Jinks-Robertson & Petes, 2021; Mateus et al., 2022). Yet, most genetic tests for sex in wild populations still assume that asyngamous recombination is negligible (e.g., Ali et al., 2016; Nicoll et al., 2024; Stoeckel et al., 2021; Tsai et al., 2008; Vakhrusheva et al., 2020), whereas others consider the evolutionary consequences of asyngamous recombination but assume obligate asexuality in the study system (e.g., Jaron et al., 2021; Jaron et al., 2022; Kershenbaum et al., 2023; Flot et al., 2013). There is hence an apparent mismatch between the bulk of theory, which assumes that asexually-produced offspring only differ by new mutations, and growing evidence that asexual reproduction involves homologous recombination.

**Fig. 1.**
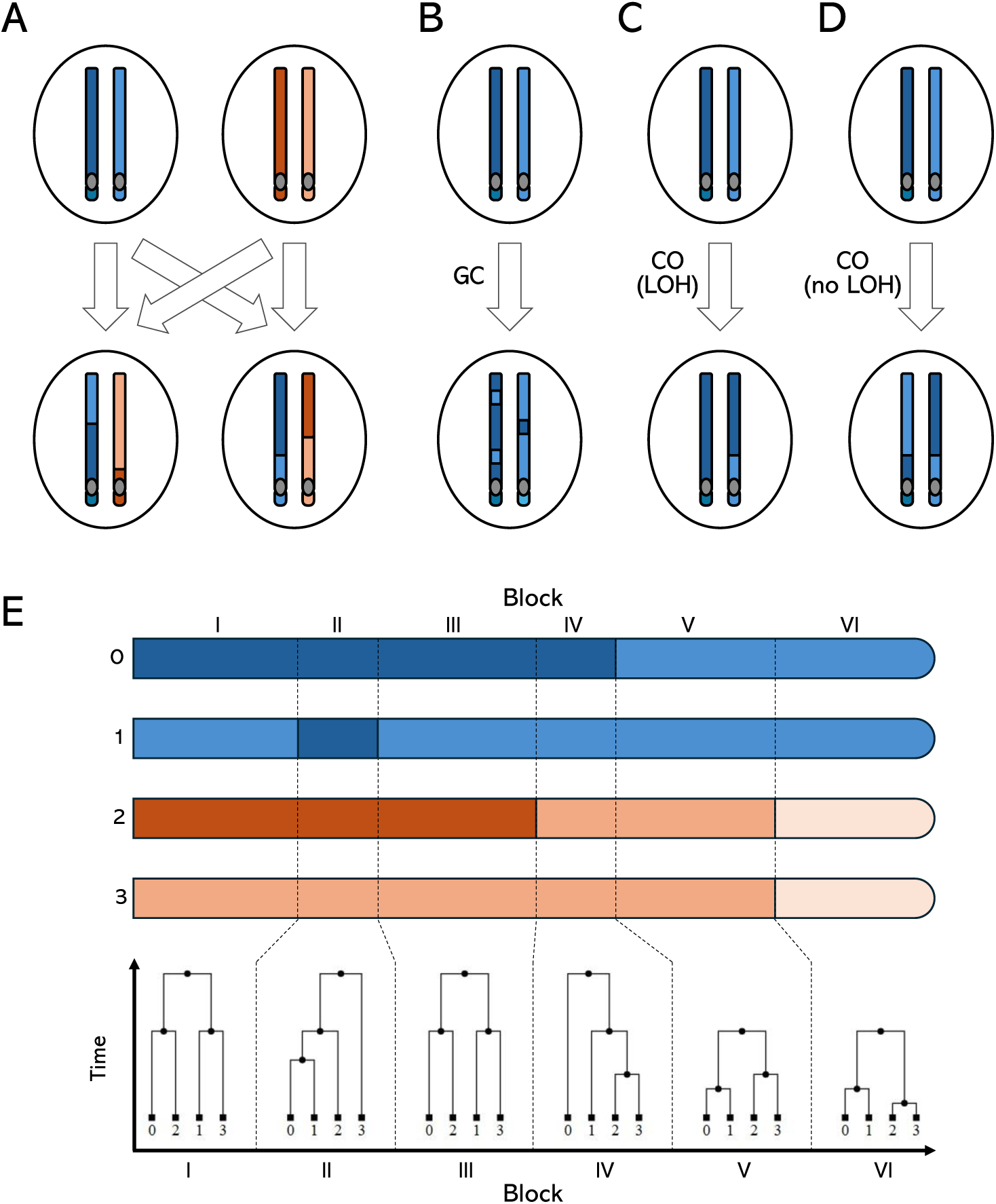
Immediate effects of sexual and asexual reproduction on sequence diversity across a diploid chromosome arm. (A) Sex with crossover recombination (left) vs asexuality (right). Asexuality can be accompanied by asyngamous gene conversion (GC; localized LOH) and crossover recombination (CO; with or without LOH). (B) Heterogenous diversity patterns reflect an underlying genealogy. In this example, the genealogy of two diploid individuals (blue and orange, with chromosomes 0, 1 and 2, 3, respectively) changes due to asyngamous recombination. The resulting patterns of diversity, with loss of heterozygosity (LOH) in one (blocks II and IV) or both (blocks V and VI) individuals, occurring more anciently (e.g., block II) or recently (e.g., block VI), are best represented as the underlying ancestral recombination graph (ARG) with local trees that describe non-recombining histories.

Another key issue is the limited consideration given to stochasticity and inference error. For instance, it has long been recognised that different regimes of DNA transmission are expected to result in distinct genealogical relationships (Birky Jr., 1996; Koufopanou et al., 1997). In diploids, clonal transmission should group homologs from different individuals as two symmetric sub-trees corresponding to allelic copies 1 and 2 (Birky Jr., 1996; Lam et al., 2011; Mark Welch & Meselson, 2000). However, past applications of these approaches have not always accounted for the various reasons why trees might deviate from the symmetric genealogy often predicted under strict asexuality. Many studies treated the discovery of any such deviations as strong evidence of genetic exchange (Debortoli et al., 2016; Laine et al., 2022; Signorovitch et al., 2015; Vakhrusheva et al., 2020; Chen et al., 2018), but inferred trees will vary for additional biological and technical reasons, including non-neutral evolution, biases in the mutation process, gene and genome duplications and loss, and reconstruction artefacts (Auxier & Bazzicalupo, 2019; Steenwyk et al., 2023; Wilson et al., 2018; Wilson et al., 2024) – alongside asyngamous recombination as outlined above.

Here, we investigate the detection and quantification of sex using individual-based, genetically explicit models. We first show that standard population genetic statistics do not distinguish between sexual and obligate asexual scenarios under realistic rates of asyngamous recombination. To overcome this challenge, we develop an analytical approach that accounts for genome-wide heterogeneity using ancestral recombination graphs (ARGs). Increasingly recognised as a natural framework to disentangle reticulate evolution (Nielsen et al., 2025; Lewanski et al., 2024; Brandt et al., 2024), ARGs describe evolutionary histories of genome regions delimited by recombination breakpoints (Wong et al., 2024) which can be represented as local (or ‘marginal’) trees (Fig. 1E). Specifically, we decompose the problem by classifying local trees for pairs of diploid individuals according to reproductive mode, and introduce ARG-inspired statistics that capture distinct signals of past reproduction. Finally, we apply our framework to a published genome dataset for the facultatively sexual budding yeast *Saccharomyces cerevisiae*, showing how novel insights can emerge from the application of ARG-based approaches to longstanding questions about reproductive mode.

## Results

### Simulation models of recombining asexuality

We constructed asexual simulation models in SLiM4 (Haller & Messer, 2023) that differ in the form of asyngamous recombination they incorporate. In the gene conversion (GC) model, representing mitotic asexuality (e.g., apomixis in animals or budding in yeast), all breakpoints initiate gene conversion tracts, each with only a local effect, that always result in LOH. In the crossover (CO) model, based on an analytical treatment of central fusion automixis (Engelstädter, 2017), recombination affects all downstream chromosomal positions, with outcomes for LOH determined by chromatid assortment and the number of crossovers per reproductive event. Full descriptions and analytical validation of both models (Engelstädter, 2017) are available in Methods and SI Appendix, section A. Unless otherwise specified, we present results based on the GC model (SI Appendix, section A highlights the conditions for equivalence versus differences between the two models).

We performed simulations using a Snakemake pipeline (Mölder et al., 2021) described in Methods and SI Appendix, section B. We simulated scenarios where a 1 Mb chromosome arm evolved neutrally in populations of *N* = 1,000 diploid individuals, reproducing sexually with probability *σ* and asexually with probability 1 - *σ*. We report the rates of sex and asyngamous recombination in terms of *N* and the asyngamous recombination rate as the per-site, per-generation rate of LOH *γ* (Kopčak & Hartfield, 2024) (SI Appendix, Table S1).

We simulated scenarios ranging from obligate asexuality (*σ* = 0), through rare sex (0 < *σ* ≤ *N^-1^*, where *N^-1^*= 10^-3^), to frequent sex (*σ* > *N^-1^*), with the rare-frequent boundary defined based on previous studies (Bengtsson, 2003; Hartfield et al., 2018). Keeping the rate of sexual crossovers fixed at a high level (10^-6^, or a single crossover per gamete on average), we varied *γ* from 0 to 5*N^-1^* (corresponding to a conversion tract initiation rate of 10^-6^ with an average tract length of 5,000 bp; Eq. 2 in SI Appendix, section A), spanning strict clonality to sex-like rates of recombination. Overall, we hence explored a two-dimensional, *σ* ∼ *γ* space of scenarios, contrasting to standard comparisons of a unidimensional clonality-to-sex continuum (Balloux et al., 2003; Halkett et al., 2005; Stoeckel et al., 2021)

### Commonly calculated statistics are often uninformative about reproductive mode

When asyngamous recombination was absent, the statistics of *H_I_* (individual-level heterozygosity), *F_IS_* (inbreeding coefficient) and *r^2^* (a measure of linkage disequilibrium decay, here calculated as an average for SNP pairs < 1 kb apart) behaved as expected from previous theory (e.g., Tibayrenc et al., 1990; Halkett et al., 2005; Stoeckel et al., 2021). Values tended towards predictions for a fully sexual, panmictic population as the rate of sex increased (Fig. 2). In the absence of sex as well as asyngamous recombination (bottom left corners), the values tended towards expectations under long-term clonality (high *H_I_* and *r^2^*, *F_IS_* ≪ 0) (Tibayrenc et al., 1990; Birky Jr., 1996). The transitions between asexuality and sex-driven values were sharpest at *σ* ≈ *N^-1^*.

**Fig. 2.**
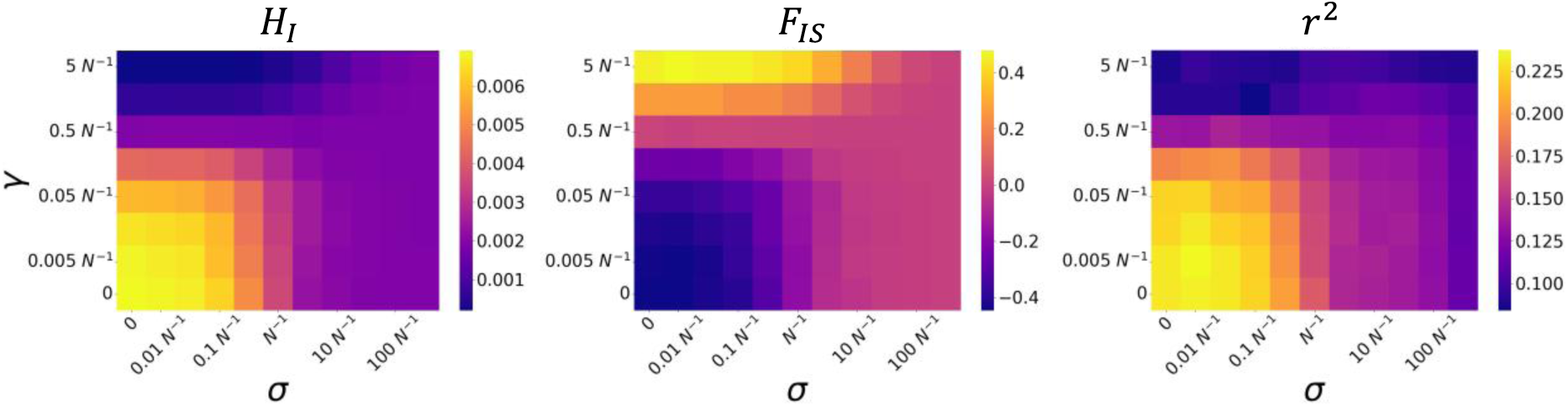
Classical population genetic statistics across the space of the sex rate *σ* and the LOH rate *γ*. For each *σ* ∼ *γ* scenario, values were computed for samples of 60 individuals and averaged across 100 replicates; *r^2^* was computed for SNP pairs < 1 kb apart. With increasing *σ*, statistic values tend towards their respective outcrossing expectations: 4*Nµ* = 0.002 for *H_I_*, 0 for *F_IS_*, and low values for *r^2^*. Note that, when *γ* ≈ 0.5*N^-1^*, ‘outcrossing-like’ statistic values are expected under obligate asexuality (*σ* = 0).

Introducing increasing rates of asyngamous recombination, however, led to a wide range of statistic values under both obligate asexuality and rare sex (Fig 2). For instance, under obligate asexuality, *H_I_* varied over six-fold across our simulation range, attaining values (<0.001–0.007) that are within the range of estimates from sexual (e.g., Mackintosh et al., 2019; Romiguier et al., 2014) and predominantly asexual groups (Jaron et al., 2021; Nowell et al., 2018). *F_IS_* varied between strongly negative values at the lower end of the simulated *γ* range (*F_IS_* ≈ -0.4) to strongly positive values at its higher end (*F_IS_* ≈ 0.4; Fig. 2), which would normally be interpreted as evidence of sexual inbreeding. Finally, average LD (SNP pairs <1 kb apart) was elevated under obligate asexuality only when *γ* < 0.5*N^-1^*, with *r^2^*= 0.23 under obligate clonality, but was otherwise low (*r^2^* ≤ 0.15) irrespective of the sex rate.

These results raise problems for a common approach to test for sex, which is to compare population genetic metrics in a target population with either a theoretical sexual expectation (e.g., Vakhrusheva et al., 2020) or patterns in related populations known to reproduce sexually (e.g., Freitas et al., 2023; Phan Thi et al., 2025), with the assumption that overlapping values are sufficient to reject obligate asexuality. On the contrary, we find that for each of the statistics, a range of LOH rates yields identical expectations for obligate asexual and frequent sex scenarios (Fig. 2). This range appears very narrow for *H_I_* and *F_IS_*, requiring *γ* of exactly 0.5*N^-1^*(Fig. 2; see also the analytical model of Vakhrusheva et al., 2020). Given a hypothetical observation of *F_IS_* = 0, we investigated whether the hypothesis of obligate asexuality could be rejected on the grounds of improbability (Vakhrusheva et al., 2020). We found that, under the GC model of asexuality, such an observation indeed requires *γ* seemingly ‘fine-tuned’ to 0.5*N^-1^*, except at very low sample size (*n* = 5; SI Appendix, Fig. S1). Under the CO model, however, a range of *γ* values (0.16*N^-1^*–1.6*N^-1^*) were likely to yield *F_IS_* = 0, even with extensive sampling (*n* = 100; SI Appendix, Fig. S1). We hence conclude that, especially in the presence of long-range recombination, observing a near-zero *F_IS_* is compatible with obligate asexuality without unrealistically stringent assumptions on the rate of asyngamous recombination.

The *γ* range where *σ* had little effect on calculated values was wider for LD decay, spanning scenarios of *γ* ≥ 0.5*N^-1^*(Fig. 2C). While recognizing that asyngamous recombination creates LD decay, some authors have interpreted certain characteristics of the decay curves as specifically indicating sex, such as the steepness of decay (Freitas et al., 2023; Vakhrusheva et al., 2020) or a specifically ‘logarithmic-exponential’ shape (Phan Thi et al., 2025) if *r^2^* is plotted as a function of inter-site distance. To evaluate these verbal arguments, we plotted *r^2^* curves across the GC and CO scenarios (SI Appendix, Figs. S2 and S3). While background LD was generally elevated under obligate asexuality, we found that asyngamous gene conversion (but not crossing-over) resulted in ‘sex-like’ shapes of decay curves (*γ* in the 0.05*N^-1^* – 0.5*N^-1^* range), with comparable magnitude of decay (drop in *r^2^* of ≈0.11 when *σ* = 0 and *γ* = 0.158*N^-1^* vs 0.15 under frequent sex) over the first 20 kb. Taken together, we conclude that the possibility of asyngamous recombination renders judgments based on classical population genetic statistics an unreliable method of detecting cryptic or rare sex.

### Tree composition analysis

We present a new method for quantitative tree-based analysis of reproductive history. This approach divides the 18 possible two-individual, four-tip coalescent trees (Wakeley, 2008) into the following categories:

**(1) Category CL: compatible with clonality (4 trees; blue in Fig. 3A).** These topologies are compatible with asexual descent without recombination (clonality) but can also be generated by other processes. In CL trees, homologs from different individuals cluster together (Birky Jr., 1996).
**(2) Category AR: compatible with asyngamous recombination (6 trees; green in Fig. 3A).** These topologies are compatible with asyngamous recombination in an asexual background. Asyngamous recombination can only result in a subset of topological changes that bring homologs together within asexual lineages (Birky Jr., 1996).
**(3) Category SX: specific to sex (8 trees; orange in Fig. 3A).** These topologies are sex-specific, i.e., are only expected through sexual reproduction or other forms of genetic exchange. SX trees are analogous to, but more general than, the ‘allele sharing’ patterns defined at the level of three diploid individuals (Laine et al., 2022; Signorovitch et al., 2015; Vakhrusheva et al., 2020; Vastrade et al., 2022).

**Fig. 3.**
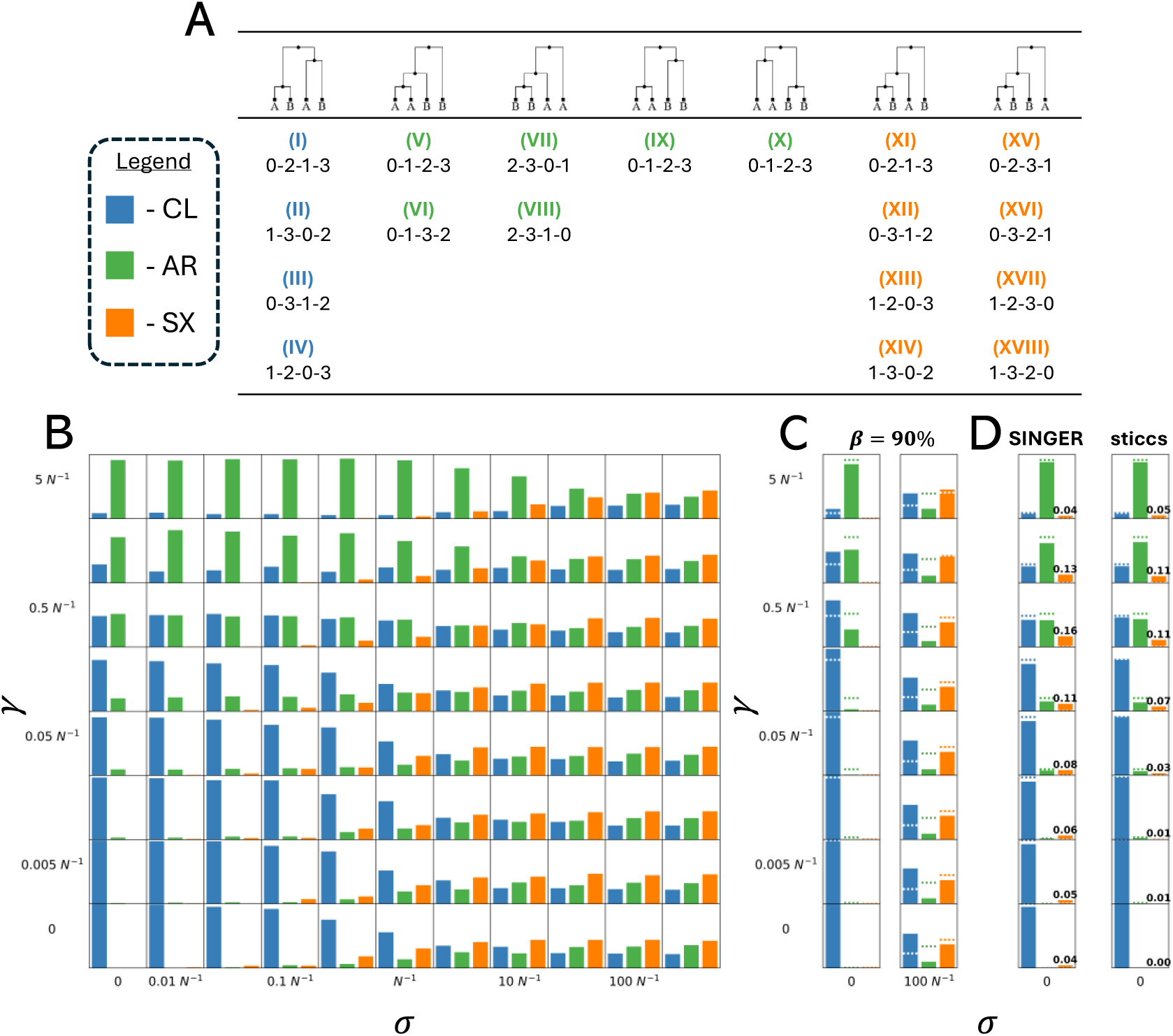
Systematic analysis of tree composition. (A) The 18 two-individual, four-tip coalescent trees can be classified into three categories: clonality-compatible (CL; blue), asyngamous recombination-compatible (AR; green) and sex-specific (SX; orange). Here, the trees are drawn for individuals A (homologs 0 and 1) and B (homologs 2 and 3), with the possible tip labelling displayed below. (B) Tree proportions *p* falling into the CL, AR and SX categories – the tree composition – across the space of the sex rate *σ* and the LOH rate *γ*. (C) A heterozygosity bias inflates *p(CL)* at the expense of *p(AR)*; here shown for two *σ* scenarios. Dashed horizontal lines represent unbiased tree composition for given scenarios. (D) Reconstruction error generates spurious SX trees; *p(SX)* reaches proportions otherwise expected under rare sex.

We propose that the contributions of asexual and sexual reproduction can be quantified as the relative genomic proportions of local CL, AR and SX trees – or a ‘tree composition’ – which takes the form of a vector (*p(CL)*, *p(AR)*, *p(SX)*). Given a genealogy for *n* individuals (2*n* tips), our approach enables exhaustive classification of signatures by aggregating them across all the possible (^*n*^) relationships (SI Appendix, Fig. S4). This pairwise decomposition moves beyond the use of trees as a categorical signature (where patterns such as ‘allele sharing’ are used as a presence-absence indicator of sex) while avoiding the combinatorial impracticality of attempting to classify all possible trees for more than two individuals. In contrast to studies that use arbitrarily delimited genome regions (e.g., entire scaffolds) to reconstruct trees (e.g., Laine et al., 2022; Öztoprak et al., 2025), an ARG framework explicitly accounts for the varying genomic span of local non-recombining trees.

### Tree composition discriminates asexual and sexual scenarios

Across the *σ* ∼ *γ* parameter space examined in our simulations, tree composition obtained by sampling two-individual ARGs reflects the rates of the underlying generative processes (Fig. 3B). Under obligate asexuality, only CL and AR trees are expected, with *p(AR)* positively correlated with rate of LOH. *p(SX)* increases with the rate of sex and, when sex is frequent, the tree composition is expected to approach a ratio of 2:3:4, reflecting the 4 CL, 6 AR and 8 SX equally probable coalescent trees. A steep transition between asexuality-like and outcrossing-like tree composition occurs at the *N^-1^* boundary between rare and frequent sex. In qualitative terms, one might distinguish three regions of the *σ* ∼ *γ* space – CL-dominant (obligate asexuality / rare sex; low LOH rate), AR-dominant (obligate asexuality / rare to frequent sex; high LOH rate), and SX-dominant (frequent sex) – approximately delimited by the *σ* = *N^-1^* and *γ* = 0.5*N^-1^* thresholds (Fig. 3B). Crucially, tree composition clearly distinguishes obligate asexual (*p(SX)* = 0) and sexual scenarios (*p(SX)* > 0), assuming that local trees can be estimated accurately.

### Heterozygosity bias in phased data exaggerates the signal of clonality

Implementing our method (or any analysis of diploid copies) requires phasing of haplotypes within individuals, but with short-read data, this is only possible for relatively heterozygous regions of the genome (Martin et al., 2016). This introduces a potential ‘heterozygosity bias’, i.e., that regions with low *H_I_* are less likely to be included. For instance, among published genomic studies on asexual animals, the proportion of the reference assembly recovered in phased segments is typically <10% (Brandt et al., 2021; Laine et al., 2022; Vakhrusheva et al., 2020) and only occasionally higher (21%; Öztoprak et al., 2025). To evaluate this issue, we added a ‘heterozygosity bias’ to our pipeline, which we denote by *β*, representing the proportion of genomic regions with insufficient *H_I_* to be sampled (see Methods). Based on estimates from real datasets (Brandt et al., 2021; Laine et al., 2022; Vakhrusheva et al., 2020), we modelled the effect of *β* = 90% across parameter space.

Adding the heterozygosity bias consistently increased *p(CL)* at the expense of *p(AR)* across the simulated parameter space (Fig. 3C; SI Appendix, Fig. S5). For example, under a frequent sex scenario of *σ* = 100*N^-1^*, *p(CL)* was inflated over two-fold relative to the outcrossing expectation (>0.5 under most values of *γ*), as would be consistent with a sex rate two orders of magnitude lower (*σ* = *N^-1^*; Fig. 3C). The effect on *p(SX)* was generally less pronounced and only occasionally led to inflated values (for some *γ* > 0.5*N^-1^* scenarios; SI Appendix, Fig. S5). Taken together, a realistic bias can drastically change the tree composition, with an overall effect of exaggerating the clonal contribution to reproductive history.

### Reconstruction artefacts introduce a spurious signal of sex

True genealogies are unknown for natural populations and must be inferred from sequence data. To investigate how reconstruction errors bias estimates of the tree composition, we performed genealogical inference on simulated datasets using two methods: SINGER (Deng et al., 2025), which identifies recombination breakpoints and infers local trees under the sequentially Markovian coalescent model (McVean & Cardin, 2005), and sticcs (Martin, 2026), which applies simple parsimony principles to the same tasks. We chose these methods as mechanistically distinct approaches with different strengths and weaknesses (see SI Appendix, Table S2 for an overview) and evaluated their performance in the absence or presence of heterozygosity bias (*β* = 90%).

Both methods erroneously inferred SX trees under obligate asexuality, with the maximum *p(SX)* of 0.164 and 0.110 for SINGER and sticcs, respectively (Fig. 3D; SI Appendix, Fig. S6). In both cases, maximum *p(SX)* occurred under *γ* = 0.5*N^-1^* – when CL and AR trees are represented approximately equally – suggesting that mixed CL-AR signal drove the mis-inference of SX trees. The better performance of sticcs under obligate asexuality can be attributed to its model-free design (Martin, 2026); on the other hand, the accuracy of SINGER increased with *σ*, consistent with an increasing match to the method’s implicit model of a sexual population (Deng et al., 2025; SI Appendix, Fig. S7). Under the heterozygosity bias, the maximum *p(SX)* under obligate asexuality was marginally lower (0.151 and 0.103 for SINGER and sticcs, respectively) and shifted to a higher *γ* of 1.58*N^-1^*(SI Appendix, Fig. S6). Taken together, we observe that two substantially different methods of genealogical reconstruction interact with asyngamous recombination to bias the tree composition, recovering SX trees at proportions otherwise consistent with rare sex when no sex, in fact, was present.

### ARG-inspired statistics to quantify signal of reproductive mode

To further quantify the signal of reproductive mode, we devised additional statistics stemming from the ARG-based framework. First, we note that the external branches of CL trees ought to have equal lengths under obligate asexuality, reflecting the number of generations to the most recent common ancestral individual. In contrast, CL trees generated through sexual exchange should exhibit unequal external branch lengths between the two subtrees, reflecting differences in the timing of internal nodes. Accordingly, our first statistic measures the degree of asymmetry in the external branch lengths of CL trees. We define *Δ* as:

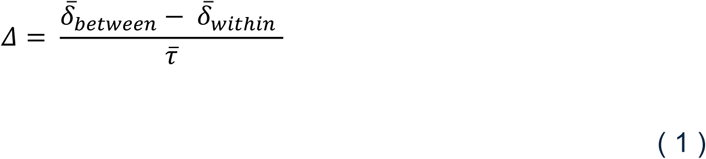

Where 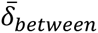 is the mean absolute difference in branch lengths *τ* between the two external subtrees, 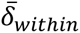 is the mean absolute difference in *τ* within subtrees, and 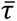 is the overall mean. An example of calculating 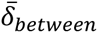 and 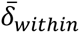 is shown in Fig. 4A. With values bounded between -1 and 2, the *Δ* statistic has an expected value of 0 given equal timing of internal coalescences, and of >0 given unequal timing within individual CL trees. The degree of deviation from 0 should be proportional to the magnitude of asymmetry in coalescence times.

**Fig. 4.**
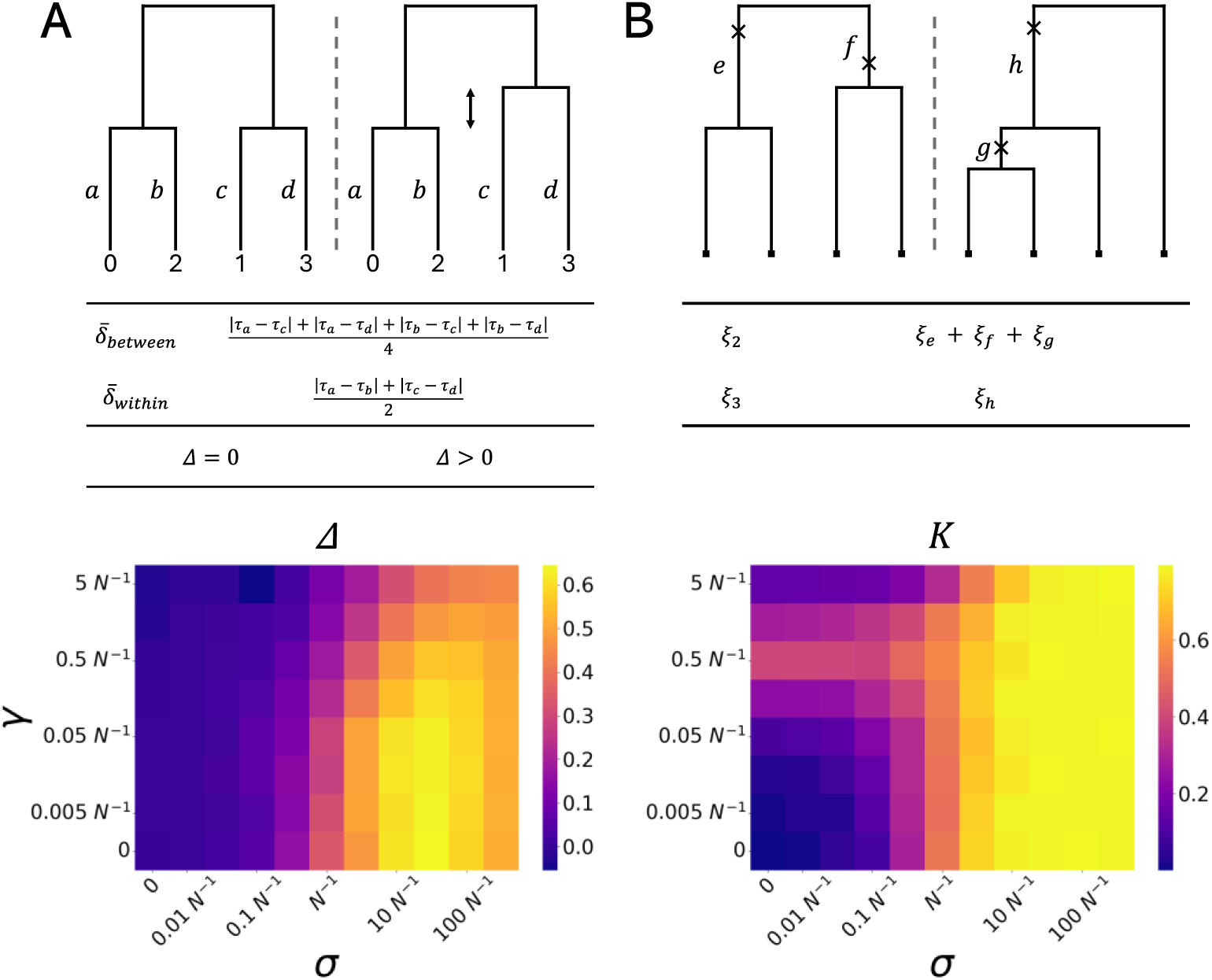
Tree-based statistics to distinguish between asexual and sexual scenarios. (A) *Δ*: a statistic calculable for haplotype-resolved segments. Top: example CL trees with equal (left) and unequal (right) timing of internal nodes. The lengths *τ* of branches *a*, *b*, *c* and *d* are used to calculate 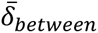 and 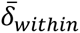; see Results. The values of *Δ* reflect the asymmetry in the timing of internal nodes in CL trees. Bottom: variation in *Δ* is driven by the rate of sex *σ*. (B) *K*: a statistic calculable for genome-wide SNPs. Top: Mutations (‘×’) arising on different branches of balanced (left; CL or AR) and unbalanced (right; AR or SX) trees contribute to the pool of doubletons (*ξ_2_*) and tripletons (*ξ_3_*). *ξ_2_* and *ξ_3_* can be used to summarise the genome-wide signal of reproductive mode; see Results. Bottom: *K* values vary with the rate of LOH *γ* but are predominantly driven by the rate of sex *σ*.

In practice, *Δ* can be computed using singleton SNPs, with the additional possibility of SNP-based delineation of target intervals (see Methods and SI Appendix, Fig. S8). In contrast to classical statistics, we found that *Δ* clearly separated obligate asexual and sexual scenarios irrespective of the asyngamous recombination rate, reaching values >0.6 under frequent sex. This discriminatory advantage of *Δ* persisted in the presence of heterozygosity bias (SI Appendix, Fig. S9). We additionally explored the distribution of *Δ* values per scenario, finding that asymmetric distributions may characterise facultatively sexual populations (SI Appendix, Fig. S10). We conclude that the *Δ* statistic captures a signal of reproductive mode that complements tree composition and remains robust under realistic assumptions of sampling bias.

Our second statistic incorporates SNPs within unphased regions to summarise the genome-wide signal of reproductive mode. We observe that, under the infinite sites model (Kimura, 1969), doubleton SNPs (with two ancestral and two derived alleles) and tripleton SNPs (with three ancestral and one derived allele per) arise on defined branches of balanced and unbalanced four-tip trees (Fig. 4B). While unbalanced 4-tip trees fall in both the AR and SX categories, they are not expected to reach a sex-like overall proportion if generated by asyngamous recombination alone (SI Appendix, Fig. S11). We define *K* (kappa) as:

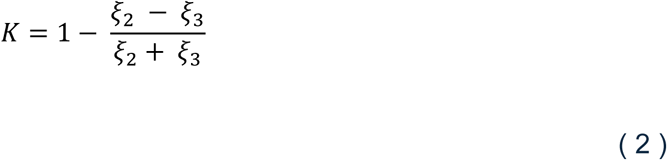

Where *ξ_2_* and *ξ_3_* are counts of doubleton and tripleton SNPs, respectively. Bounded between 0 and 2, the *K* statistic has an expected value of ≈ 0.79 under frequent sex. Under obligate asexuality, the expectation varies with the asyngamous recombination rate but does not exceed ≈ 0.39 (Fig. 4B). Expected values that do not overlap between the opposite ends of the asexuality-sex spectrum set *K* apart from the standard summary statistics of unphased sequence variation (cf. Fig. 3). Finally, to incorporate variation across simulation runs, we used a simple approximate Bayesian computation (ABC) routine (SI Appendix, Fig. S12), finding that *Δ* and *K* consistently outperformed classical statistics in predicting *σ*.

### Genealogical analysis reveals contrasting reproductive histories across *S. cerevisiae* clades

To assess the potential of our framework to uncover novel biological insights, we applied it to four clades of the budding yeast *Saccharomyces cerevisiae* (Peter et al., 2018) chosen to exemplify variation in expected life cycles (Becerra-Rodríguez et al., 2026). The life cycle of *S. cerevisiae* alternates between frequent diploid, mitotic reproduction and rarer meiotic production of haploid spores. Ploidy can be restored either by fusion with a product of the same meiotic division – via intra-tetrad mating (Nishant et al., 2010) or haplo-selfing (Haber, 2012) – or by fusion with an unrelated spore (outcrossing). Based on laboratory assays, Becerra-Rodríguez et al. (2026) assigned *S. cerevisiae* isolates to three broad life cycle types. The ‘conventional’ life cycle involves all three mechanisms of ploidy restoration; ‘preferred sexual’ is defined by defective haplo-selfing and a presumed preference for outcrossing; finally, ‘preferred asexual’ is defined by defective sporulation (complete failure or inviable progeny) and putatively obligate mitotic reproduction. If *S. cerevisiae* clades do follow such distinct life cycles in nature, we predict an excess of AR trees for ‘conventional’ clades (assuming a major role of LOH-inducing haplo-selfing), SX trees for ‘preferred sexual’ clades 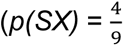, if sex is frequent), and CL trees for ‘preferred asexual’ clades (with *Δ* = 0 and *p(AR)* dependent on the rate of asyngamous recombination).

Leveraging telomere-to-telomere, haplotype-resolved genome assemblies (Loegler et al., 2025), we performed genome-wide genealogical analyses of target clades (Methods). We uncovered marked variation in reproductive histories among clades, sometimes confirming and sometimes contradicting expectations derived from the laboratory assay (Fig. 5; see SI Appendix, section C for a detailed account). Confirming expectations, the West African Cocoa clade, with all isolates classified as ‘conventional’, exhibited the expected AR-dominant tree composition at the clade level (*p(AR)* = 0.406), albeit with considerable variation in genealogical signatures among pairs of isolates (SI Appendix, Fig. S13).

**Fig. 5.**
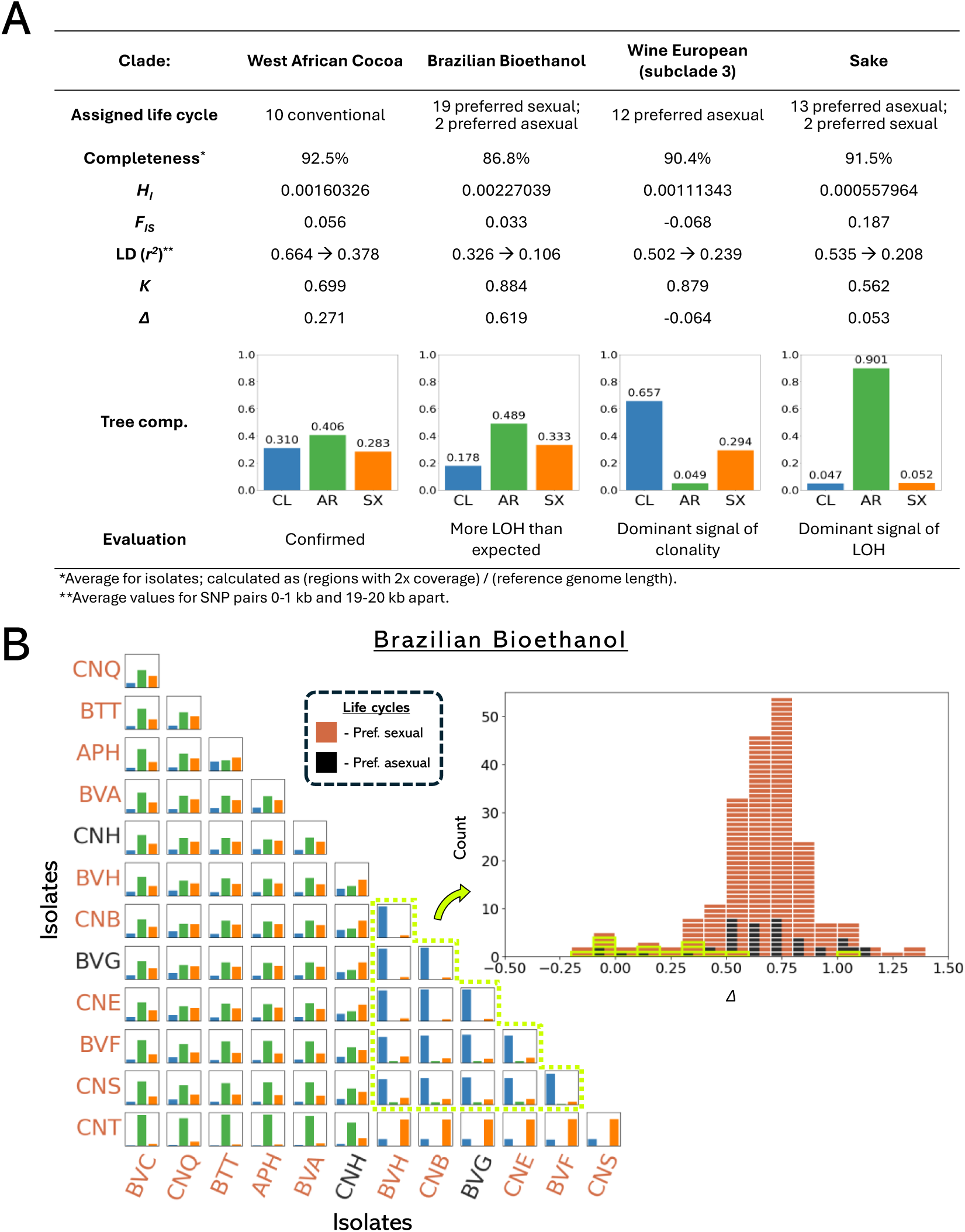
Reproductive signatures in four clades of *Saccharomyces cerevisiae*. (A) Evaluation of life cycle assignments of Becerra-Rodríguez et al. (2026) based on classical and genealogical metrics. (B) Example of further analysis based on the genealogical framework: the Brazilian Bioethanol clade (see also SI Appendix, Figs. S13 and Fig. S14 for the remaining clades). Isolate codes follow Loegler et al. (2025). Analysis of pairwise tree composition (left) and the distribution of *Δ* values (right) identifies a putative asexual subclade (yellow), characterized by a CL-dominant tree composition and a distinct, 0-overlapping distribution of *Δ* values.

Contradicting expectations, while classified as largely ‘preferred sexual’, the Brazilian Bioethanol clade displayed an even higher proportion of AR trees (*p(AR)* = 0.489), suggesting a non-trivial role of LOH-inducing processes. This result coincided with the identification of a distinctive CL-dominant, and presumably asexual, sub-clade (Fig. 5B; see SI Appendix, Figs. S13 and S14 for another putative example in West African Cocoa). While both the Wine European and Sake clades consisted predominantly of ‘preferred asexual’ isolates, an assignment supported by near-0 values of *Δ* (-0.064 and 0.053, respectively), they appeared to differ markedly in the proportion of AR trees and hence rates of asyngamous recombination, which was substantially higher in the Sake clade (Fig. 5A). Importantly, these patterns were not readily evident from classical statistics (SI Appendix, section C), which provided an incomplete (*F_IS_*) or ambiguous picture (*H_I_*, LD decay). Further exploration of genealogical signatures, including across chromosomes (SI Appendix, Fig. S15), could provide previously inaccessible insights into the evolutionary ecology of *S. cerevisiae*, and non-model reproductive systems more broadly.

## Discussion

Estimating rates of sexual and asexual reproduction is a prerequisite for answering a range of fundamental evolutionary questions. Interpreting genomic variation through the lens of reproductive history could shed light on deep links between reproductive mode and microevolutionary patterns (Feretzaki & Heitman, 2013; Morran et al., 2011), macroevolutionary trends (Barraclough, 2019), and a range of ecological, genetic, and life history peculiarities (Jeffries et al., 2025; Wilson et al., 2024). While jointly modelling multiple forces that shape genome-wide variation has become a gold standard across many branches of evolutionary research (Laetsch et al., 2023; Johri et al., 2022; Excoffier et al., 2021), this approach has yet to be widely adopted for inferring reproductive histories. As a step toward addressing this issue, we focus on homologous recombination that operates independently of the sexual pathway, which remains largely overlooked in current approaches to quantifying sex from sequence data (e.g., Ali et al., 2016; Nicoll et al., 2024; Stoeckel et al., 2021; Tsai et al., 2008; Vakhrusheva et al., 2020).

Using simulation models of asexuality, we found that asyngamous recombination confounds sequence-based statistics commonly used to support the presence of sex (e.g., Freitas et al., 2023; Phan Thi et al., 2025; Vakhrusheva et al., 2020). Specifically, these statistics can take values characteristic of outcrossing populations even when sex is rare or absent. In many asexual systems, estimates of the per-site rate of LOH are consistently on the order of 10^-5^-10^-4^ (Table S3), broadly aligning with the levels required to mimic sex-like patterns of variation (for *N* on the order of 10^-3^-10^-4^). Although classical population-genetic summaries do retain some information about reproductive mode – for instance, all else being equal, background LD should be elevated under obligate asexuality (SI Appendix, Figs. S2 and S3) – their use to support reproductive hypotheses has typically relied on assumptions that recombination is ‘rare’ and exclusively ‘short-range’ (Berry et al., 2019; Koester et al., 2018; Lasek-Nesselquist et al., 2009; Rogers et al., 2014; Van den Broeck et al., 2020; Weedall et al., 2012; Phan Thi et al., 2025). Our analysis of an *S. cerevisaie* population genomic dataset further demonstrated that LOH levels can vary widely across asexual clades of the same species, or even within such clades, suggesting sensitivity of the underlying mechanisms to ecological and genetic contexts that warrants further investigation.

Although it might seem improbable that asyngamous recombination would take exactly those values required to confound tests of sex, there are biological reasons why this might be expected. The generation of LOH has two contrasting consequences for fitness. On the one hand, exposure of recessive deleterious mutations leads to loss of fitness and inbreeding depression (Charlesworth & Willis, 2009). On the other, LOH can help fix beneficial mutations and counteract Muller’s ratchet by overwriting deleterious mutations (Flot et al., 2013; Kopčak & Hartfield, 2024; Mandegar & Otto, 2007). To maximise fitness benefits, selection may affect rates of spontaneous asyngamous recombination and LOH in a similar way to recombination rates in sexual organisms (Dapper & Payseur, 2017), or via programmed distortion of chromatid segregation (Blanc et al., 2023; Lacy et al., 2024). Concurrently, observed levels of LOH might be biased towards intermediate values through selection against excess homozygosity (Flynn et al., 2017).

The majority of ARG reconstruction methods are rooted in a simple model of a sexual population with crossover recombination (Li & Stephens, 2003; McVean & Cardin, 2005). Here, we show that ARG-based approaches can be extended to systems with non-canonical reproductive biology. Using *S. cerevisiae* as a case study with recent high-quality datasets (Becerra-Rodríguez et al., 2026; Loegler et al., 2025), we demonstrated how novel biological insights can arise from such applications. Most strikingly, we found that laboratory assays of reproductive mode (Becerra-Rodríguez et al., 2026) can bear little correspondence to long-term reproductive history, which we interrogated at the isolate level through pairwise comparisons. Even with haplotype-resolved assemblies, we nonetheless caution that limitations to inference itself can still bias the biological insights. For example, while the high overall *p(SX)* in the Wine European clade could be interpreted as evidence against its obligate asexuality, other mechanisms generating tripleton SNPs (e.g., homoplasy) might produce a similar excess. Development of genealogical methods that explicitly model non-canonical reproductive modes will be critical for resolving ambiguities of this kind.

As advances in long-read sequencing make genome-wide genealogical inference increasingly feasible across non-model systems, simulation models offer a powerful foundation for statistical inference using approximate Bayesian computation (Beaumont, 2010), supervised machine learning (Schrider & Kern, 2018), and convolutional neural networks (Whitehouse et al., 2024; Korfmann et al., 2023). One evolutionary conundrum that could benefit from these new datasets and approaches is the contentious status of so-called ‘ancient asexuals’ (Judson & Normark, 1996), with claims of cryptic sex in the well-documented absence of males (Laine et al., 2022; Vakhrusheva et al., 2020; Wilson et al., 2024; Birky Jr, 2010), and conversely, claims of long-term asexuality in their presence (Öztoprak et al., 2025; Taberly, 1988). We suggest that resolution can be achieved by (1) explicitly accounting for asyngamous recombination, (2) addressing technical and methodological limitations, and (3) quantifying reproductive mode signatures within a statistical framework. We provide the building blocks for this aim.

## Methods

We used tskit v0.5.8 (Kelleher et al., 2016; Wong et al., 2024) to process simulated ARGs, scikit-allel v1.3.13 (Miles et al., 2024) to work with SNP data, and matplotlib v3.9.2 (Hunter, 2007) and seaborn v0.13.2 (Waskom, 2021) for visualisation in Python. The simulation pipeline and analysis code are available at https://github.com/TymekPieszko/SexSigns.

### Simulation models of asexuality

We implemented both the GC and CO models in scenarios where a single diploid chromosome of length *L* evolved in populations conforming to Wright-Fisher assumptions (discrete, non-overlapping generations, constant population size), in which offspring are produced sexually by hermaphroditic parents with probability *σ* and asexually with probability 1 - *σ* (see SI Appendix, Table S1 for definitions of all simulation parameters). Asexual reproduction is accompanied by asyngamous recombination: gene conversion with an initiation rate *r_GC_* in the GC model and crossover recombination at rate *r_CO_* in the CO model (see Supplementary Text 1 for descriptions of the recombination routines). The length of gene conversion tracts is geometrically distributed with mean *λ* and can be set independently of the tract initiation rate. Sexual reproduction is accompanied by crossover recombination at rate *r* using the standard machinery of SLiM4.

We validated both models against analytical predictions for equilibrium individual-level heterozygosity *Ĥ*_*I*_ (Engelstädter, 2017; see SI Appendix, section A for a summary). For the CO model, consistency with the analytical model was obtained by assuming that crossovers occur between all four chromatids, i.e., in the absence of chromatid interference (SI Appendix, Fig. S16). For both models, we obtained close concordance between analytical and simulated values (*N* = 100 individuals, 5 × 10^5^ generations, 30 replicates per scenario, other parameters as in SI Appendix, Table S1), except at low recombination rates under the CO model (*γ* ≤ 0.05*N^-1^*; SI Appendix, Fig. S17), where this runtime was still insufficient to reach mutation-LOH equilibrium. Although finite runtimes necessarily limit how closely individual-based simulations can match analytical expectations in such edge cases (including in our main pipeline; see below), equilibria are unlikely to be encountered in nature for the same reason.

### Simulation pipeline

We ran our simulations in a pipeline written using the workflow management system Snakemake v8.13.0 (Mölder et al., 2021). In summary, the pipeline consisted of three major steps. First, we ran simulations with tree-sequence recording of output (Haller et al., 2019) across the space of sex rates *σ* and LOH rates *γ* (see SI Appendix, Table S1 for all parameter values used). Each scenario was run for 5,000 generations and replicated 100 times. Second, after ensuring complete coalescence of simulated ARGs using pyslim v1.0.4 (Haller et al., 2019), we subsampled each ARG to two random individuals and generated files for downstream processing of these target samples. Using msprime v1.3.2 (Baumdicker et al., 2022), we added neutral mutations at a per-site per-generation rate *µ* = 5 × 10^-7^ chosen to avoid the computational burden of simulating larger populations (SI Appendix, section B). Finally, for each two-individual ARG, we re-inferred genealogies using sticcs v0.0.5 (Martin, 2026) and SINGER v0.1.8-beta (Deng et al., 2025). Details of the pipeline are described in SI Appendix, section B.

### Calculating population genetic statistics

We calculated classical statistics, as well as *K* and *Δ* (see below), using sub-ARGs of 60 random individuals, processed analogously to the two-individual ARGs generated within the simulation pipeline. We calculated *H_I_* using a custom function, *F_IS_* using scikit-allel, and *r^2^* for SNP pairs < 1 kb apart using tskit’s ‘LdCalculator’. For heatmap visualisation, we calculated mean values for each statistic across 100 replicates within each σ ∼ γ scenario. To visualize LD decay curves, we used 10-individual sub-ARGs and 100 randomly sampled focal SNPs, binning *r^2^* values by distance in 1-kb windows.

### Tree composition analysis

We wrote a tree-classification function using the ‘tracked_samples’ functionality in tskit, which we applied to both simulated and inferred ARGs. We applied the heterozygosity bias post hoc, by dividing the simulated chromosomes into windows of 5 kb, ranking them in by increasing *H_I_*, and sampling the highest-ranking windows. To obtain tree composition from SINGER output, we performed additional averaging across samples of the MCMC chain.

### New genealogical statistics

We provide Python functions for calculating *K* and *Δ*, with two alternative approaches available for the latter. The first uses CL-tree intervals defined from a simulated or reconstructed genealogy; for example, in the *S. cerevisiae* analysis, we used intervals inferred by sticcs. The second, illustrated in SI Appendix, Fig. S8, bypasses genealogical reconstruction by approximating CL-tree locations directly from the configurations of doubleton SNPs. Across the *σ* ∼ *γ* space of simulated scenarios, this SNP-based procedure accurately reproduced *Δ* values obtained from the true CL intervals (SI Appendix, Fig. S9). In both approaches, the counts of the four singleton-SNP classes within each interval serve as proxies for the lengths of the corresponding external branches.

### *S. cerevisiae* analysis

We restricted the analysis to isolates classified as both diploid and euploid (Table S4; Peter et al., 2018). For each of 58 such isolates, we aligned the two haplotype assemblies (66% of which consisted of 16 haplotypes corresponding one-to-one with haploid chromosomes) against the S288C reference genome using minimap2 v2.17 with the asm5 preset (Li, 2018). We extracted variant sites from each haplotype-to-reference alignment using paftools.js, and merged the resulting haploid VCF files into single-isolate diploid VCFs using bcftools merge and bespoke code. Finally, we merged the per-isolate sets of variants into a single file, retaining 26,270 biallelic SNPs for downstream processing. All the ‘bcftools merge’ operations were run with the ‘--missing-to-ref’ flag.

We polarised SNPs using a divergent *S. cerevisiae* clade from Taiwan (Peter et al., 2018). We cleaned and filtered Illumina paired-end reads for the three available Taiwanese isolates (Peter et al., 2018) using ‘bbduk’, performed error correction using ‘tadpole’ from the BBTools v37.62 suite (Bushnell, 2017), and aligned them against the reference using minimap2 v2.17 (Li, 2018). We called variants using ‘bcftools call’, retaining biallelic SNPs with QUAL ≥ 30. As a polarisation rule, we assumed that ≥ 5 alternative alleles in the outgroup alignment provide evidence for re-polarisation of a site, ≤ 1 asserts correct polarity of a site, with the remaining sites ambiguous and hence excluded. To obtain an analysis-ready set of SNPs, we implemented per-isolate masking of alignment gaps (sites with <2x coverage per isolate) and centromeric regions, the latter defined using the S288C annotation. Per-isolate ‘callable’ sequence covered between 87% and 93% of the reference genome across the four clades.

We computed population-level values of classical and genealogical statistics, weighing estimates by chromosome length and, for *Δ*, by the span of CL intervals inferred for each pair of isolates (see below). We inferred ARGs for all (^*n*^) pairs of target isolates per clade using sticcs v0.0.5 (Martin, 2026) with ‘forced_ploidy = 1’ (as required for running the algorithm on haplotype-resolved data) and ‘second_chances = True’. ARG inference was restricted to regions jointly ‘callable’ for both isolates per pair, leaving edge-free gaps in the resulting tree sequences.

## Acknowledgments

We thank Dmitri Filatov, Aziz Aboobaker, Alan Grafen, Antoine Houtain and Reuben Nowell for comments on earlier versions of the work and manuscript drafts. Simulations were run on the University of Oxford Advanced Research Computing (ARC) facility (http://dx.doi.org/10.5281/zenodo.22558). T.P. was supported by the UKRI Natural Environment Research Council and Clarendon Fund Scholarships. J.K. was funded by the Robertson Foundation. T.G.B. and C.G.W. were funded by UKRI Natural Environment Research Council grants NE/M01651X/1 and NE/S010866/2.

