## Supplementary material for "Detecting and quantifying rare sex in natural populations": SI Appendix

### Supporting Information Text

#### A: Models of asexuality with asyngamous recombination

We validated our models against analytical predictions for equilibrium individual-level heterozygosity $\widehat{H}_I$. For the CO model, these predictions are based on Engelstädter (2017), who considered automixis via central fusion (Stenberg and Saura, 2009). While the terms 'automixis' and 'central fusion' imply a specific cytological pathway (see Blanc et al., 2025, for a recent critique), the CO model applies more generally to forms of asexuality involving crossover recombination and co-segregation of non-sister chromatids (e.g., Beukeboom & Pijnacker, 2000; Eisman & Kaufman, 2007; Hiruta et al., 2010; Terwagne et al., 2022). Below, we summarise the analytical expectations and explore alternative design choices for the CO model. We then describe our implementation in an individual-based SLiM4 framework (Haller & Messer, 2023) and finally compare the GC and CO models.

Analytical expectations

During asexual reproduction, heterozygous genotypes are either passed intact to offspring or transition to a homozygous state through asyngamous recombination. Engelstädter (2017) provides a general formula for $\widehat{H}_I$ under obligate asexual reproduction:

$$\widehat{H_I} \approx \frac{2\mu}{2\mu + \gamma} \tag{1}$$

Where $\mu$ is the per-site per-generation mutation rate and $\gamma$ is the probability of a transition from the heterozygous state in the parent to a homozygous state in the offspring (i.e., the rate of loss of heterozygosity specific to a site), with the approximation valid for small $\mu$. Under mitotic asexuality with gene conversion, an expression for $\gamma$ can be obtained simply, following Engelstädter (2017):

$$\bar{\gamma}_{GC} = r_{GC} \times \lambda$$

( 2 )

Where $r_{GC}$ is the rate of gene conversion tract initiation and $\lambda$ is the mean gene conversion tract length. Together, Eq. (2) and Eq. (1) provide an expectation for equilibrium individual-level heterozygosity, against which we compare our GC model (Fig. S17).

Implementing asyngamous crossing-over in an individual-based framework requires additional design choices. For example, to simplify the model, one may assume that all crossovers occur exclusively between a single pair of chromatids. This assumption corresponds to complete negative chromatid interference (Sarens et al., 2021; hereafter, the 'CI' assumption) and has been used in previous simulation work (Blanc et al., 2023). Under the CI assumption, crossovers between the centromere and a focal locus induce switches between 2 configurations: recombinant (for odd crossover numbers) and non-recombinant (for even crossover numbers). Given *n* crossovers between the centromere and a focal locus, the probability of arriving at the recombinant state *r(n)* is:

$$r(n) \mid CI = \sum_{i=1}^{n} (-1)^{i-1}$$
$$= \frac{1}{2}(1 - (-1)^n)$$

( 3 )

If the number of crossovers is Poisson-distributed with mean $\bar{n}$, Eq. (3) can be modified to obtain the expected fraction of recombinant configurations at a locus using Taylor series expansion:

$$\bar{r}(\bar{n}) \mid CI = \sum_{k=0}^{\infty} \frac{\bar{n}^k \, e^{-\bar{n}}}{k!} r(k)$$

$$= \frac{1}{2}\sum_{k=1}^{\infty}\frac{\bar{n}^k\, e^{-\bar{n}}}{k!}(1-(-1)^n)$$

$$= \frac{1}{2}\left(1\ -\ e^{-\bar{n}}\ -\ \sum_{k=1}^{\infty}\frac{\bar{n}^k\, e^{-\bar{n}}}{k!}\right)$$

$$= \frac{1}{2}\left(1-e^{-\bar{n}}-e^{-\bar{n}}(e^{-\bar{n}}-1)\right)$$

$$= \frac{1}{2}(1-e^{-2\bar{n}})$$

( 4 )

Finally, by incorporating the co-segregation of non-sister chromatids (for a given crossover, a recombinant and a non-recombinant chromatid have a 50% chance of segregating together; see Fig. 1 in Engelstädter, 2017), we obtain $\gamma_{CO}$, the probability of a transition from the heterozygous state in the parent to a homozygous state in the offspring:

$$\gamma_{CO}\ \Big|\ CI\ =\ \frac{1}{2}\ \bar{r}(\bar{n})\ \Big|\ CI$$

$$= \frac{1}{4}(1-e^{-2\bar{n}})$$

( 5 )

This can be contrasted with the expression derived by Engelstädter (2017) under the assumption that crossovers occur with equal probability between all four possible pairs of non-sister chromatids, corresponding to no chromatid interference (‘NCI’):

$$\gamma_{CO}\mid NCI\ =\ \frac{1}{3}\left(1-e^{-\frac{3\bar{n}}{2}}\right)$$

( 6 )

In Fig. S18, we compare the values of $\widehat{H}_I$ obtained by combining Eq. (1) with Eq. (5) or Eq. (6) across a range of expected crossover numbers $\bar{n}$. As expected, the match is close when $\bar{n} \leq 0.01$, but decays with higher values of $\bar{n}$. In other words, the CI assumption provides a good approximation to Engelstädter's (2017) analytical model only at low $\bar{n}$, whereas the NCI routine (Fig. S16) is required for full consistency. It remains unclear which assumption, if either, better reflects the biology of asexual groups (Blanc et al., 2025).

Finally, the mean chromosome-wide rate of LOH under the CO model $\bar{\gamma}_{CO}$ can be obtained as:

$$\bar{\gamma}_{CO} = \frac{1}{L-1}\sum_{i=1}^{L-1}\gamma_i \tag{7}$$

Where $L$ is the chromosome length in number of sites.

<u>Model description</u>

SLiM4 (Haller & Messer, 2023) is a scriptable framework for constructing individual-based, genetically explicit simulations. While SLiM4 supports both sex and clonality, it does not natively support asyngamous recombination. A workaround is to draw breakpoint positions using custom code in SLiM's Eidos scripting language and generate offspring 'genomes' (SLiM's term for single DNA molecules) using the genomes of a single parent as templates. Below, we outline the steps for drawing recombination breakpoints under the GC and CO models, as well as for incorporating sexual and asexual reproduction in simulated scenarios.

**Gene conversion in the GC model.** The generation of offspring genomes is accompanied by a Poisson-distributed number of gene conversion tracts with mean $\bar{n}_{GC}$. Each tract, with mean length $\lambda$, is modelled as two breakpoints marking its start and end. The Poisson distribution of tract number is achieved by following a Poisson process with interarrival times (here, distances between the end of one tract and the start point of the next) geometrically distributed with parameter $r_{GC}$; hence, $\bar{n}_{GC}$ = $r_{GC}$ × $L$, assuming $\lambda \ll L$. Implementation of the Poisson process specific to the GC model is outlined in Algorithm A; note that the algorithm does not avoid edge effects.

**Algorithm A**. Legend: $i$ – current position, $i_{max}$ – maximum position, $D$ – distance to next breakpoint, $i_{start}$ – track start position, $i_{end}$ – tract end position, $Z$ – length of a tract (see also Table S1).

| | |
|---|---|
| | Set $i$ = 0, $i_{max}$ = $L$ and BREAKS = [] |
| | **while** $i$ < $i_{max}$ **do** |
| | Draw $D$ = rand(Geom($r_{GC}$)) |
| | Set $i_{start}$ = $i$ + $D$ |
| | **if** $i_{start}$ > $i_{max}$ **then** |
| | **exit** |
| | Draw $Z$ = rand(Geom($\frac{1}{\lambda}$)) |
| | Set $i_{end}$ = $i_{start}$ + $Z$ |
| | **if** $i_{end}$ > $i_{max}$ **then** |
| | **exit** |
| | BREAKS.append([$i_{start}$, $i_{end}$]) |
| | Set $i$ = $i_{end}$ |
| **Return** | BREAKS |

Conversion tracts drawn via Algorithm A are randomly assigned to the four possible pairs of non-sister chromatids. Finally, parent genomes are randomly assigned to each of the two ‘slots’ of the offspring individual (i.e., passed in random order to the addRecombinant() method), and offspring genomes incorporating the conversion tracts are generated.

**Crossover recombination in the CO model.** The generation of offspring genomes is accompanied by a Poisson-distributed number of crossovers with mean $\bar{n}_{CO}$. Each crossover is modelled as a breakpoint that simultaneously affects two chromatids, resulting in a switch in the parental copying template. The Poisson distribution of crossover number is achieved by following a Poisson process with interarrival times (here, distances between breakpoints) geometrically distributed with parameter $r_{CO}$; hence, $\bar{n}_{CO}$ = $r_{CO}$ × $L$ (Table S1).

Implementation of the Poisson process specific to the CO model is outlined in Algorithm B; note that the algorithm does not avoid edge effects.

**Algorithm B.** Legend: $i$ – current position, $i_{max}$ – maximum position, $D$ – distance to next breakpoint, $i_{CO}$ – position of a crossover (see also Table S1).

Set $i$ = 0, $i_{max}$ = $L$ and BREAKS = []
**while** $i$ < $i_{max}$ **do**
    Draw $D$ = rand(Geom($r_{CO}$))
    Set $i_{CO}$ = $i$ + $D$
    **if** $i_{CO}$ > $i_{max}$ **then**
        **exit**
    BREAKS.append($i_{CO}$)
    Set $i$ = $i_{CO}$
**Return** BREAKS

Crossovers drawn via Algorithm B are first randomly assigned to the four possible pairs of non-sister chromatids. These assignments are then modified to reflect how the physical cutting and rejoining of DNA molecules affects the outcomes at linked loci, a process of 'crossover resolution' illustrated in Fig. S16. Finally, parent genomes are randomly assigned to each of the two 'slots' of the offspring individual (i.e., passed in random order to the addRecombinant() method), and offspring genomes are generated according to the resolved crossovers.

**Mixed reproductive scenarios.** We followed section 15.13 of the SLiM4 manual (Haller & Messer, 2024) to achieve Wright-Fisher-like dynamics using the non-Wright-Fisher machinery of SLiM4. Briefly, generation $t$ + 1 is produced by sampling generation $t$ with replacement, with sampling probabilities proportional to fitness (equal for all individuals under neutral scenarios). In our mixed reproductive scenarios, two pools of parents are sampled. If an offspring is to be produced asexually, a parent from pool 1 is used; if an offspring is to be produced sexually, a parent from pool 1 is paired with a parent from pool 2.

Model comparison

**Expected values.** As for the GC model, we simulated scenarios across the $\sigma \sim \gamma$ range discussed in the Main Text using the CO model. To parameterize CO simulations appropriately, we used optimisation via Brent's method implemented in scipy v1.17.0 (Virtanen et al., 2020), such that for each value of $\gamma$ we obtained the corresponding value of $r_{CO}$ (Table S1). For the different calculated metrics, the mean chromosome-wide values matched closely between the two models (e.g., Fig. A). Genealogies simulated under CO nonetheless displayed a heightened *p(AR)* (Fig. B), which we attribute to stronger edge effects (see 'Model description' above). Overall, we conclude that focusing on the GC model was a justifiable choice.

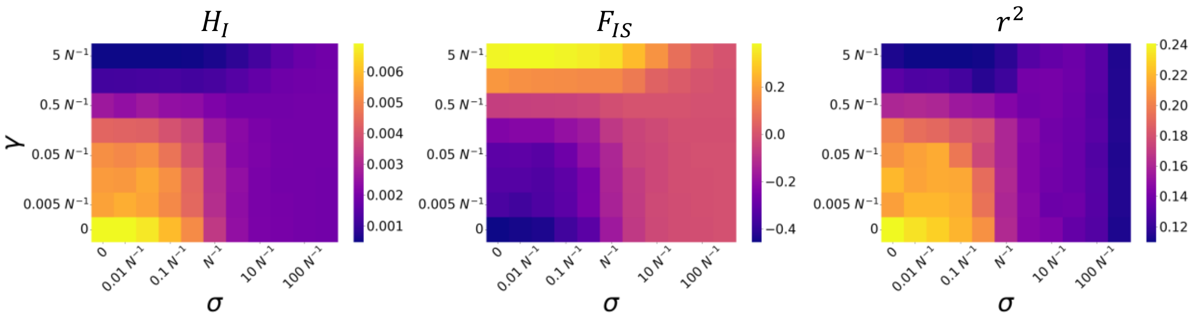


**Fig. A.** Classical population genetic statistics across the space of the sex rate $\sigma$ and the LOH rate $\gamma$: CO model (cf. Fig. 2, Main Text). Statistic values were computed as described in the legend to Fig. 2, Main Text.

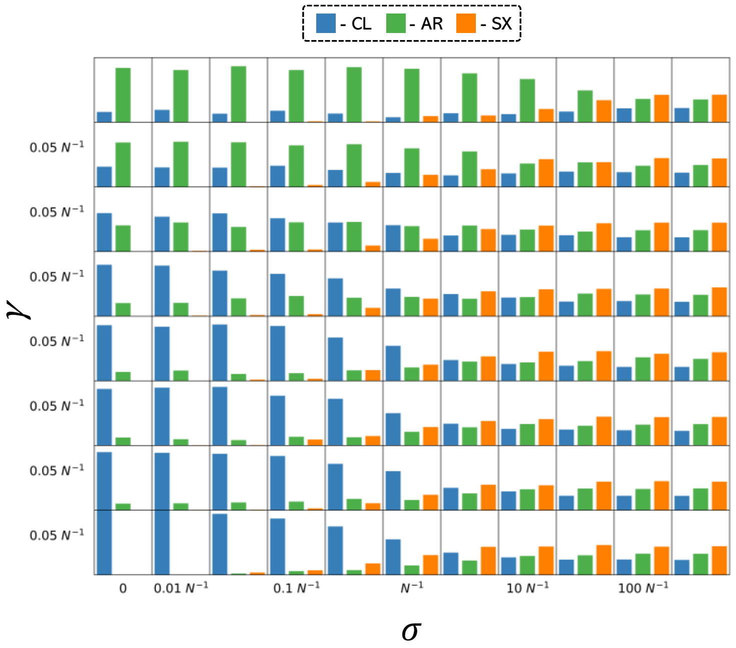


**Fig. B.** Tree composition across the space of the sex rate $\sigma$ and the LOH rate $\gamma$: CO model (cf. Fig. 3B, Main Text).

**Chromosomal heterogeneity.** The GC and CO models differed in the distribution of diversity across chromosomes: uniform in the GC model but decaying from position 0 to position $L$ in the CO model (Fig. S17). The pattern associated with the CO model reflects our assumption that position 0 is the 'centromere' and position $L$ is the 'telomere', as in one arm of a monocentric chromosome. Alternative designs could instead represent multiple centromeres distributed uniformly across the sequence, as in holocentric chromosomes (Blanc et al., 2023). These considerations are not central to our present aims but may become important when modelling a specific asexual system.

**Variance.** As evident for $F_{IS}$ from Fig. S1, CO model was associated with greater variance of computed values across replicates of each scenario. This can be understood by noting that the average number of bases affected by each recombination event is larger in the CO than in the GC model: accordingly, $r_{CO}$ values were approx. two orders of magnitude lower than $r_{GC}$ for equivalent values of $\gamma$ (Table S1). A related result is the relative flatness of LD decay curves under CO (compare Figs. S2 and S3). While considerations of variance were important to several analyses presented here – including the two examples above and the ABC analysis – we note that differences between these outcomes should be interpreted relatively rather than absolutely, with chromosome length as a confounding factor.

Taken together, the present study does not fully explore the differences between GC and CO models, leaving room for future investigations. At the same time, it is worth noting that, while theoretical analyses treat asyngamous gene conversion and crossing-over separately (e.g., Kopčak & Hartfield, 2024; Engelstädter, 2017), such 'pure' scenarios are unlikely to occur in nature. Indeed, sequencing studies typically report the co-occurrence of LOH tracts across multiple genomic scales (e.g., Vijayan et al., 2025; Dutta et al., 2021; Houtain, Derzelle, Llirós, et al., 2024).

### B: Simulation pipeline

Below, we provide details on the processing of simulated ARGs and the re-inference of genealogies implemented in the Snakemake pipeline (Mölder et al., 2021) used to run our simulations.

Processing of simulated ARGs

We performed several in-memory operations on the simulated tree sequences. First, to ensure complete coalescence of the genealogies, we performed recapitation using pyslim v1.0.4 (Haller et al., 2019). 'Recapitation' is a post-hoc procedure of adding a sexual burn-in for a hypothetical sexual progenitor of the asexual lineage using a coalescent simulator msprime (Baumdicker et al., 2022), for which we used an ancestral $N_a$ = 1000 and crossover rate $r_a$ = $10^{-6}$. Second, we subsampled each simulated ARG to two random individuals. We then used msprime v1.3.2 (Baumdicker et al., 2022) to overlay neutral mutations under the Jukes-Cantor model at a per-site per-generation rate $\mu = 5 \times 10^{-7}$ on top of the subsampled ARGs. This value of $\mu$ is two orders of magnitude higher than is typically reported in invertebrates (Wang and Obbard, 2023); paired with $N$ = 1,000, it was chosen to avoid the computational burden of simulating very large populations. Under the traditional scaling approach (Uricchio and Hernandez, 2014), these parameter values would approximate the evolutionary processes in a population of 100,000 individuals. We nonetheless note that our non-canonical models are unlikely to conform to the scaling trick and, accordingly, we report the simulated rates of $\gamma$. Per simulation, we output the two-individual ARG (for downstream tree-based and SNP-based steps) and a corresponding VCF file (input for SINGER).

Re-inference of genealogies

Using both sticcs and SINGER, we re-inferred genealogies for the target two-individual ARGs. These contained 436 to 5,490 non-recombining blocks (Dataset 1, Table S5) and 3,205 to 8,046 SNPs on average (Dataset 1, Table S6) in GC scenarios, allowing us to test the inference algorithms' performance in the presence of mixed genealogical signal. We ran sticcs v0.0.5 (Martin, 2026) with 'forced_ploidy = 1' (as required for running the algorithm on haplotype-resolved data) and 'second_chances = True'. For SINGER re-inference, we used the two-individual VCF files as input. As scaling parameters, we used the simulated values of $N$ and $\mu$. We kept the recombination **:** mutation rate ratio of 1 across the simulated scenarios and set '-polar 0.99' (i.e., SNPs are correctly polarized). Per run of the MCMC chain, we output 10 samples, evenly distributed between steps 1,000 and 2,000 of the chain. Although the SINGER algorithm terminates when the mismatch between the input diversity and the expectation of $4N\mu$ exceeds a certain threshold, we did not observe any failures across the simulated scenarios.

### C: *S. cerevisiae* analysis

With a thorough mechanistic understanding of the life cycle (Haber, 2012; Börner et al., 2023) and the molecular pathways of recombination (Symington et al., 2014), the budding yeast *Saccharomyces cerevisiae* is an ideal target for investigating reproductive mode evolution in natural, or domesticated, populations. Most reproductive events in the *S. cerevisiae* life cycle are mitotic, with estimates of the rate of spontaneous LOH on the order of $10^{-5}$ per bp per generation (Table S3). This rate is a resultant of diverse molecular pathways (e.g., homologous recombination or non-homologous end-joining) generating a diversity of LOH patterns, including interstitial (from < 100 bp to megabase-scale LOH in the middle of chromosomes) and terminal tracts (generally kilobase to megabase-scale LOH at chromosome ends) (Vijayan et al., 2025; Dutta et al., 2021). Following meiotic sporulation, which produces a tetrad of haploid spores enclosed in an ascus, ploidy is restored through the fusion of spores of opposite mating types – **a** and α – encoded at the *MAT* locus (Haber, 2012). Several different pathways of ploidy restoration are available. Intra-tetrad mating involves the fusion of *MAT***a** and *MAT***α** spores of the same tetrad**.** The so-called 'haplo-selfing' occurs when a mitotic division of a haploid spore is followed by the parent cell switching its mating type and fusing with its daughter cell (Haber, 2012). Finally, outcrossing occurs when spores are released from the ascus and, following mitotic divisions in the environment, encounter a compatible spore. Using strategies that ignore mitotic recombination (Ruderfer et al., 2006; Tsai et al., 2008) or, conversely, use LOH events to date the timing of past outcrossing (Magwene et al., 2011), it has been previously estimated that sporulation in *S. cerevisiae* and its sister species *S. paradoxus* occurs once per $10^3$-$10^5$ mitotic generations. Intriguingly, more recent estimates based on large genomic datasets are, to our knowledge, lacking.

The sequencing of over 1,000 *S. cerevisiae* genomes (Peter et al., 2018; Loegler et al., 2025) has provided an invaluable genomic resource for evolutionary studies. A recent laboratory assay (Becerra-Rodríguez et al., 2026) scored 670 of these isolates for major components of the sexual life cycle: the ability to enter meiosis and sporulate under starvation, the viability of meiotic progeny following tetrad dissection, and the capacity of haploid spore progeny to undergo mating-type switching. The authors proposed that the *S.*

*cerevisiae* life cycle can be broadly divided into three types (Becerra-Rodríguez et al., 2026), as outlined in Main Text. Before considering the correspondence between population-genetic signatures and assigned life cycles, several general points are worth noting. First, although elevated $H_I$ is commonly regarded a signal of outcrossing (Peter et al., 2018), we exclude it from this discussion because high $H_I$ can arise through both outcrossing and long-term clonality. Although LOH rates suggest that the accumulation of a clonal signature should generally be limited (Table S3), our results indicate that this may not always be the case (Wine European; see Fig. 5 and below). Indeed, even a comparison of $H_I$ and $F_{IS}$ values for West African Cocoa and Brazilian Bioethanol clades shows that $H_I$ (Fig. 5A) is not a reliable indicator of reproductive mode. We also note that LD decay was observed in all clades, as expected under both sexual and asyngamous forms of recombination. Disentangling the relative contributions of these processes (Hartfield et al., 2018) within an explicit inference framework is beyond the scope of this study.

A second point is that, despite near-chromosome-level phasing, the phased genomic dataset (Loegler et al., 2025) represents only a subset of a larger genomic dataset (Peter et al., 2018). The isolates selected for phasing were not chosen at random and, by necessity, excluded homozygous genomes (Table S4). We therefore flag, on a clade-by-clade basis, cases in which this ascertainment may influence interpretation.

West African Cocoa clade (10 isolates, 45 pairwise comparisons)

All isolates of the West African Cocoa clade were classified as possessing the 'conventional' life cycle and could therefore be expected to be largely homozygous as a consequence of haplo-selfing. Seemingly consistent with this expectation, the clade displayed the highest level of background LD ($r^2$ = 0.378 vs. $r^2$ ≤ 0.239 in other clades). A positive but near-zero $F_{IS}$ = 0.056 indicated, however, that asexuality and / or outcrossing played a non-trivial role. Specifically, the presence of frequent outcrossing ($\sigma > N^{-1}$) was supported by $K$ = 0.699, indicating an excess of unbalanced topologies beyond that expected from within-lineage

LOH alone. This latter result was reflected in a tree composition of $c$ = (0.310, 0.406, 0.283), nonetheless displaying an excess of CL and AR trees over the outcrossing expectation. While overall $\Delta$ = 0.271 suggested imbalance within CL trees providing additional evidence for genetic exchanges, a subclade of isolate pairs with CL-dominant tree composition showed $\Delta$ values enriched in the 0-overlapping tail of the distribution (Figs. S13 and S14), suggesting asexual descent. Taken together, these results are only tentatively consistent with the assigned life cycle type, with clearer evidence for outcrossed and asexual groups of isolates rather than highly selfing lineages. Importantly, this conclusion is not biased by the availability of phased genomes, since 10 / 13 isolates from the original dataset (Peter et al., 2018) were included in the phased dataset (Loegler et al., 2025). With all classified as 'heterozygous', we assume the choice of isolates was haphazard.

Brazilian Bioethanol clade (21 isolates, 210 pairwise comparisons)

The majority (19 / 21) of isolates from the Brazilian Bioethanol clade were classified as 'preferred outcrossing', with the remaining two classified as 'preferred asexual'. In agreement with this classification, the calculated statistics ($F_{IS}$ = 0.033, background $r^2$ =0.106, $K$ = 0.884, $\Delta$ = 0.619) were consistent with a largely panmictic population. The overall tree composition of $c$ = (0.178, 0.489, 0.333) nonetheless displayed an excess of AR trees over the outcrossing expectation; this included 168 / 210 AR-dominant pairs of isolates (*p(AR)* = 0.571). On the other hand, 6 isolates displayed a strikingly CL dominant tree composition (*p(CL)* = 0.847), with $\Delta$ values enriched in the 0-overlapping tail of the distribution (Main Text, Fig. 5B), suggesting a clonal subclade. Interestingly, only one of these isolates was originally classified as 'preferred asexual'. Similarly to the West African Cocoa clade, these findings are only tentatively consistent with the assigned life cycle type, with heightened levels of LOH in some isolates and asexuality in others. Importantly, the observed *p(AR)* is likely an underestimate of true clade-level value, with 26 / 35 of isolates classified as 'heterozygous', pushing the clade further from outcrossing expectations.

Wine European clade (12 isolates, 66 pairwise ARGs)

All isolates of the Wine European clade were classified as possessing the 'preferred asexual' life cycle. Despite an only mildly negative, near-zero $F_{IS}$ = -0.068, and K = 879, the overall tree composition was CL-dominant: *c* = (0.657, 0.049, 0.294). A near-zero $\Delta$ = -0.064 provided additional support for a largely asexual history and the assay-based classification. At the same time, the relatively high *p(SX)* (cf. Fig. 3D) admits two possible interpretations. First, the overall proportion could in principle indicate a degree of outcrossing, contrasting sporulation failure in laboratory conditions. However, examination of the distribution of *p(SX)* values across pairwise comparisons (Fig. S13) revealed very similar levels among pairs, suggesting that much of this signal may instead reflect a consistent background source of error, such as homoplasy or polarisation error. The original set of Wine European isolates consisted of 'heterozygous' isolates only (24 / 24), with a presumably haphazard selection of genomes for phasing.

Sake clade (15 isolates, 105 pairwise ARGs)

Similarly to the Wine European clade, the majority (13 / 15) of Sake isolates were classified as possessing the 'preferred asexual' life cycle. The patterns in the Sake clade nonetheless stood in stark contrast to Wine European. A positive $F_{IS}$ = 0.178 was reflected in an AR-dominant tree composition of *c* = (0.047, 0.901, 0.052), with 87 / 105 AR-dominant pairs of isolates (*p(AR)* = 0.943), and the remaining pairs showing more evenly distributed tree composition suggestive of outcrossing. Intriguingly, $\Delta$ = 0.053, although this estimate should be interpreted cautiously given the low overall proportion of CL trees. Another surprising result was $K$ = 0.562, seemingly inconsistent with both the observed *p(AR)* – corresponding to an expectation of $K \approx 0.145$ under obligate asexuality – and the exceptionally low *p(SX)*. One possibility is a systematic difference between balanced and unbalanced AR trees, but further investigation should exclude an artefactual source of surplus tripleton SNPs. Despite these caveats, the overall pattern appears consistent with predominantly asexual reproduction accompanied by a very high rate of LOH. Accordingly, 38 / 47 isolates from the original dataset were ‘heterozygous’, with the remaining ones likely to contribute similar or even higher *p(AR)*.

### Supporting Information Figures

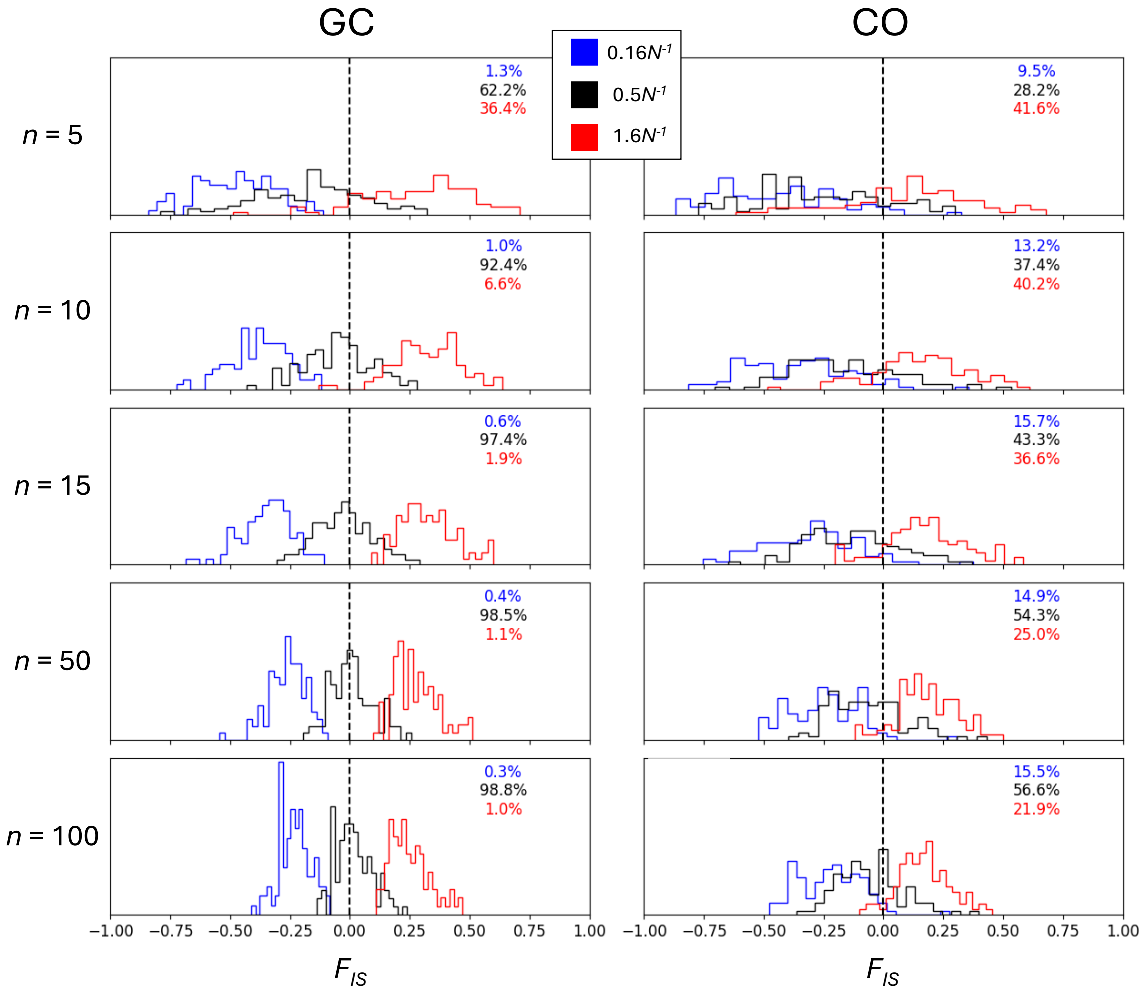


**Fig. S1.** Likelihood of observing $F_{IS}$ = 0 under obligate asexuality, sample sizes (*n*) between 5 and 100, and different values of *γ*; comparison between the GC and CO models. Likelihoods of $F_{IS}$ = 0 were obtained assuming normal distribution of simulated values, transformed to Akaike information criterion (AIC) values (with one parameter), and finally Akaike weights of evidence for different values of *γ* (top right of each panel).

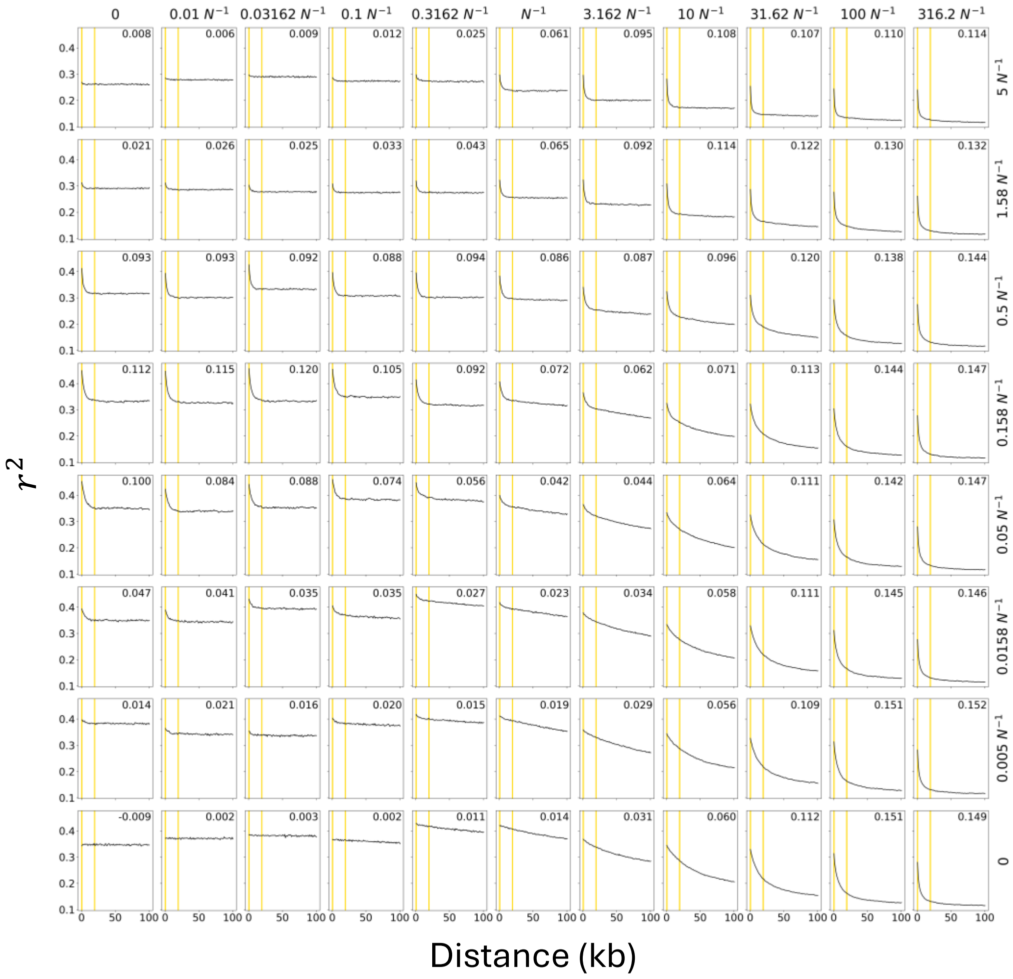


**Fig. S2.** $r^2$ as a function of inter-SNP distance across the space of the sex rate $\sigma$ and the LOH rate $\gamma$.; GC model. $r^2$ was calculated using samples of 10 individuals and 100 focal SNPs per simulation.

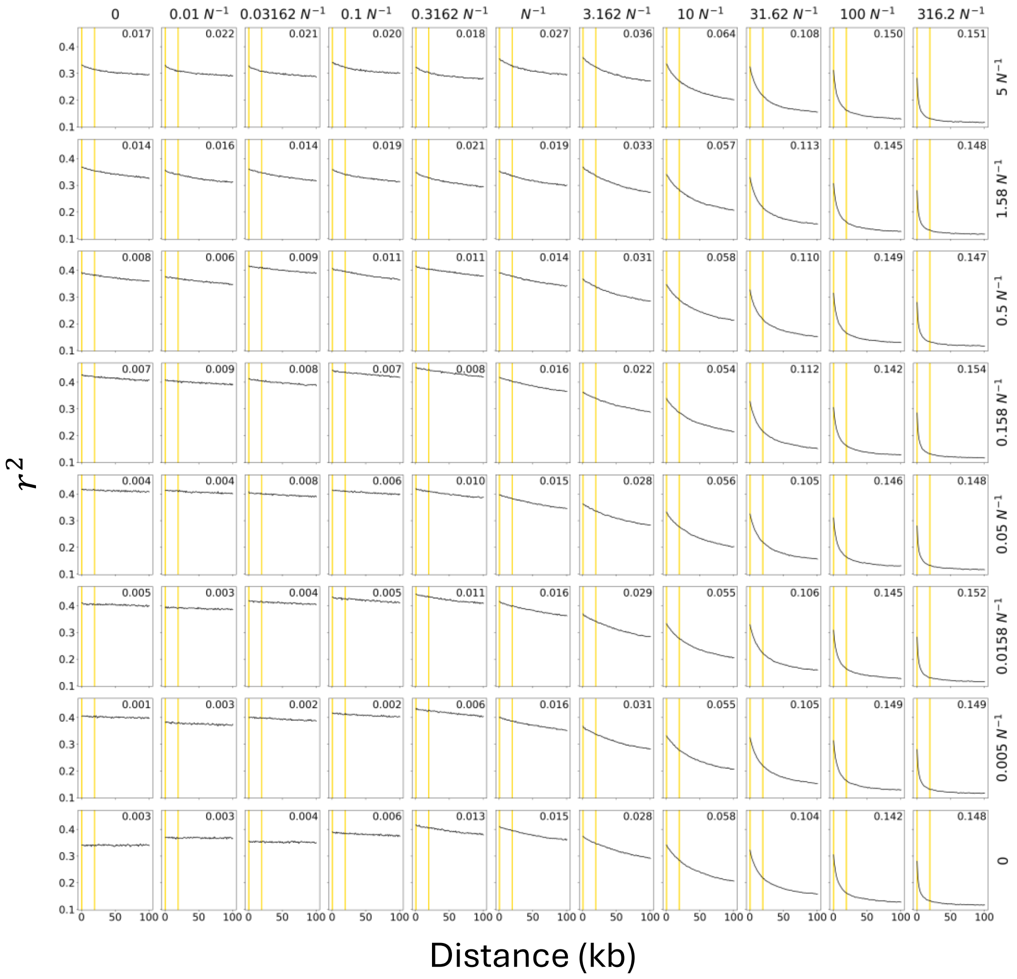


**Fig. S3.** $r^2$ as a function of inter-SNP distance across the space of the sex rate $\sigma$ and the LOH rate $\gamma$.; CO model. $r^2$ was calculated using samples of 10 individuals and 100 focal SNPs per simulation.

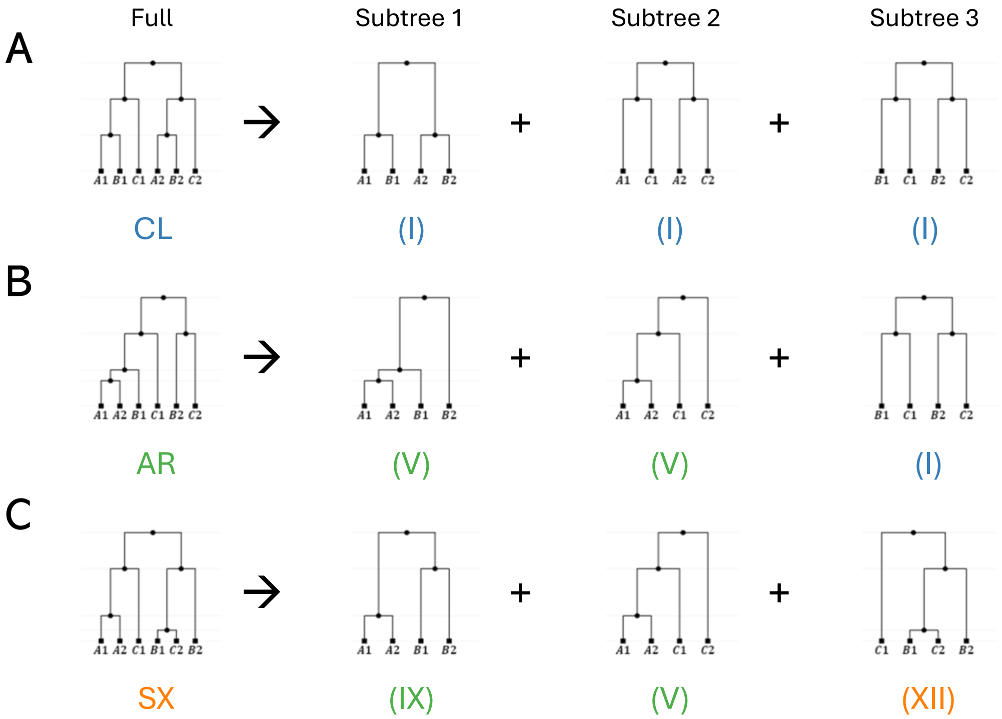


**Fig. S4.** Exhaustive tree classification: example of 3 individuals. To disentangle the reproductive history, trees for individuals $\boldsymbol{A}$, $\boldsymbol{B}$ and $\boldsymbol{C}$ (each with alleles 1 and 2) can be decomposed into subtrees for the $\binom{3}{2}$ possible pairs of individuals. Roman numerals and colours correspond to Fig. 3A. (A) A clonal tree is decomposed into CL subtrees (note the equal timing of internal nodes that is not recovered by many inference algorithms and hence ignored in Fig. 3A). (B) The signature of asyngamous recombination is reflected in subtrees 1 and 2. (C) The signature of sex is reflected in subtree 3.

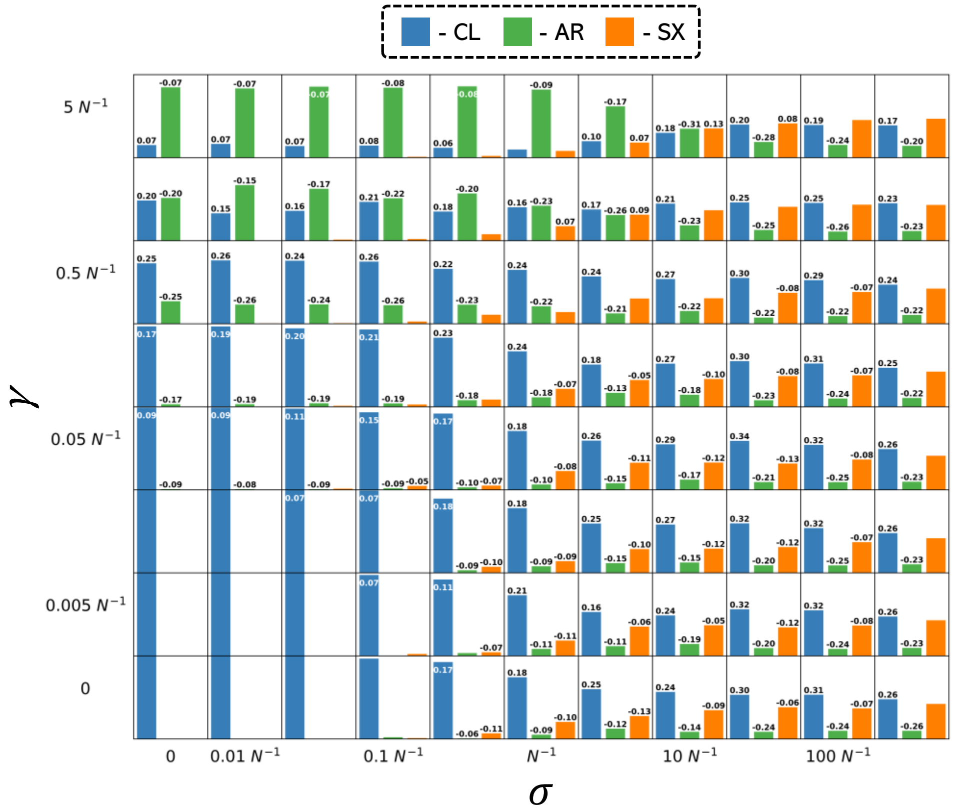


**Fig. S5.** Tree composition across the space of the sex rate $\sigma$ and the LOH rate $\gamma$; $\beta$ = 90%. Differences from the simulated ground truth (no bias) are annotated if they exceeded 0.05.

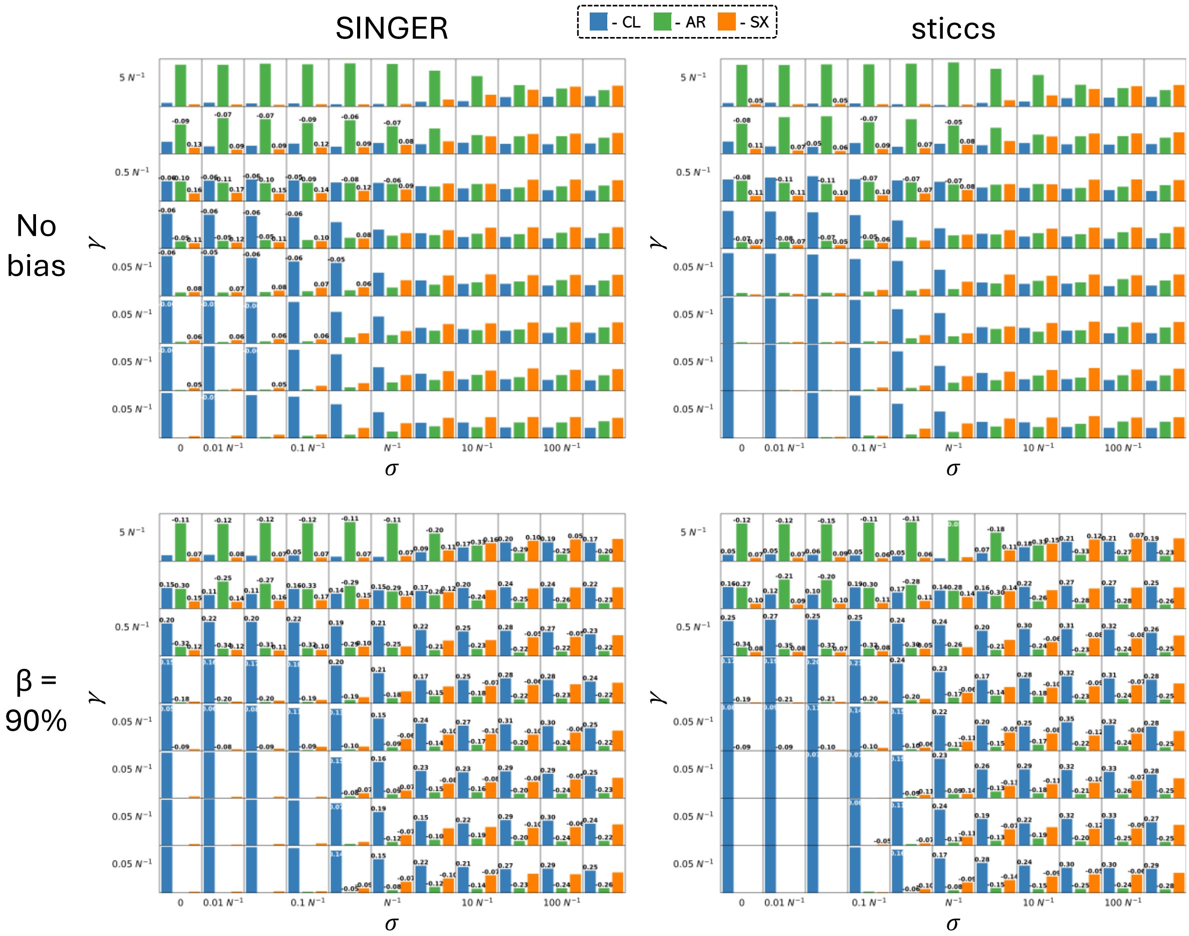


**Fig. S6.** Tree composition across the space of the sex rate $\sigma$ and the LOH rate $\gamma$; reconstruction with SINGER and sticcs. Differences from the simulated ground truth (no reconstruction, no bias) are annotated if they exceeded 0.05.

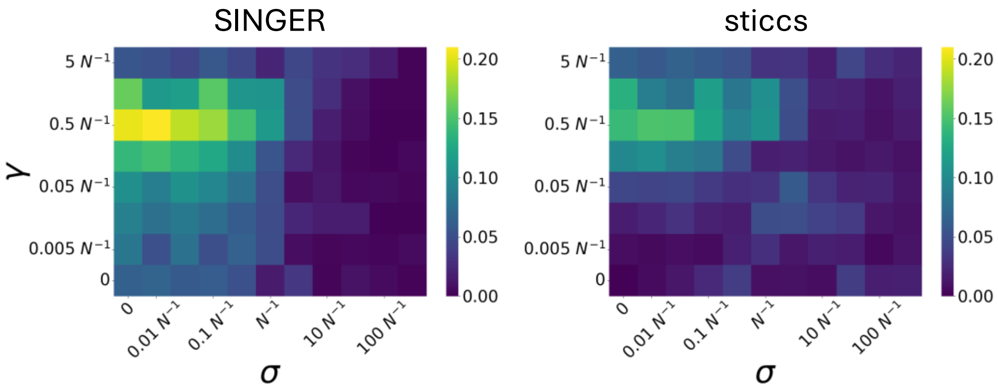


**Fig. S7.** Tree composition: Euclidean distance from simulated ground truth across the space of the sex rate $\sigma$ and the LOH rate $\gamma$ and after reconstruction with SINGER and sticcs.

(A)

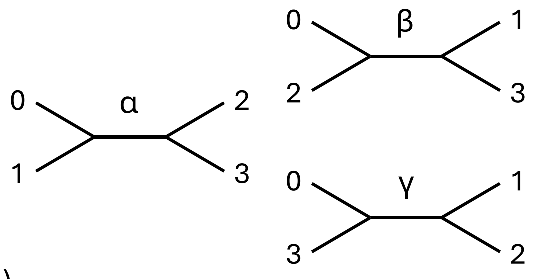


| Topology | Haplotype 0 | 1 | 2 | 3 |
|---|---|---|---|---|
| α | - | - | × | × |
| | × | × | - | - |
| β | - | × | - | × |
| | × | - | × | - |
| γ | - | × | × | - |
| | × | - | - | × |

(B)

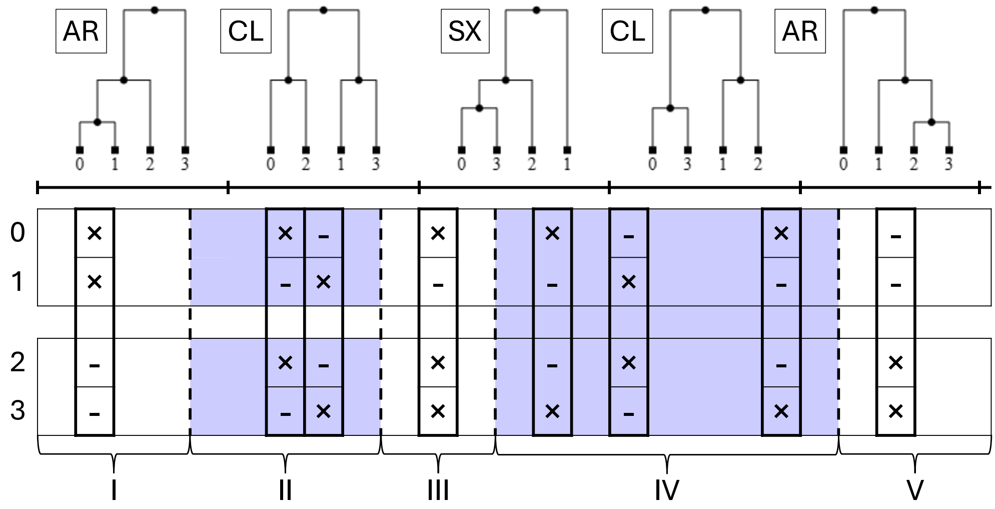


**Fig. S8.** An SNP-based procedure for calculating *Δ*. (A) The three topologies of unrooted trees with four tips (left). Under the infinite sites model, pairs of doubleton SNP classes (right; derived alleles denoted with '×') correspond uniquely to one of the unrooted topologies. (B) Local four-tip trees are reflected in the runs of doubleton and tripleton SNPs. The intervals of CL trees, all of which are balanced, can be approximated as unbroken runs of doubleton SNPs corresponding to unrooted trees β and γ (here, intervals II and IV, respectively). Note that, although depicted as single trees in the schematic, runs are expected to span multiple local trees sharing the same topology. A tripleton indicates an underlying unbalanced rooted topology (interval III). For computing *Δ*, intervals are delimited by mid-points between terminal doubletons of target SNP runs, and singletons (now shown) are used as a proxy for the length of external branches.

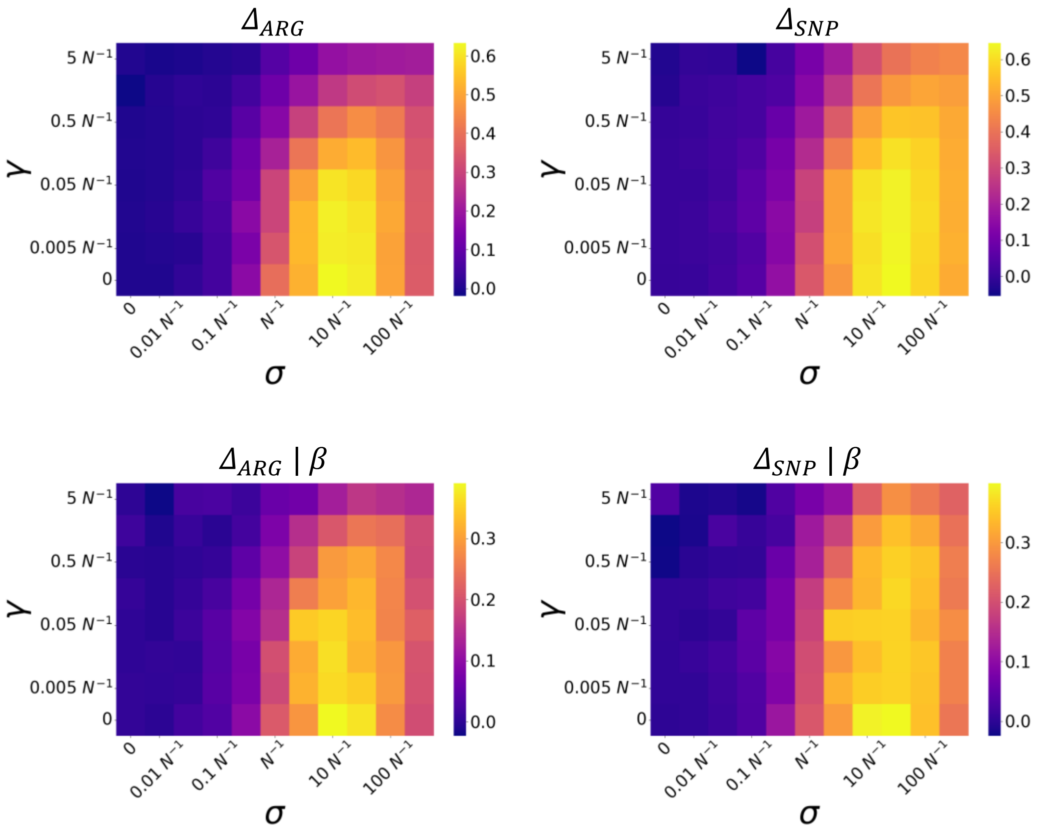


**Fig. S9.** $\Delta$ across the space of the sex rate $\sigma$ and the LOH rate $\gamma$. $\Delta_{ARG}$ and $\Delta_{SNP}$ denote the ARG-based and SNP-based procedures, respectively, of delimiting CL intervals outlined in Methods. For $\Delta_{ARG}$, true simulated ARGs were used; β = 90%.

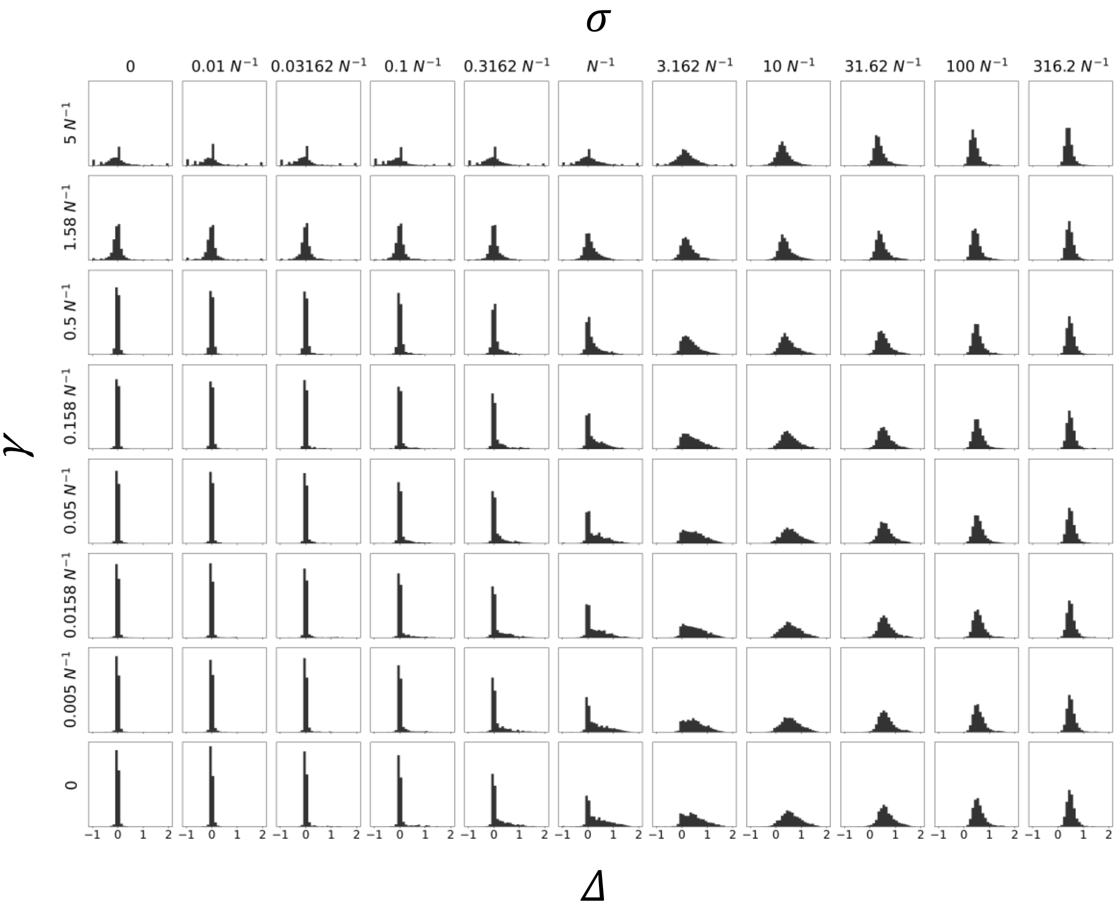


**Fig. S10.** Histograms of *Δ* values under scenarios across the space of the sex rate *σ* and the LOH rate *γ*; GC model. Each histogram consists of 3,000 values (100 replicates × 30 pairs of individuals per replicate).

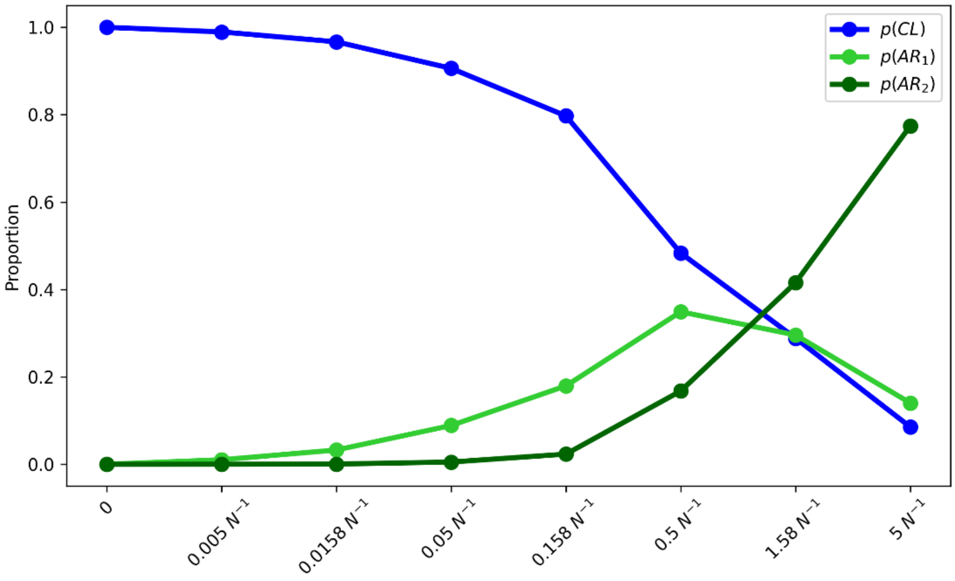


**Fig. S11.** Proportions of CL, unbalanced AR ($AR_1$) and balanced AR ($AR_2$) trees across the range of $\gamma$ values under obligate asexuality; GC model. Here, the maximum *p*($AR_1$) reached is approx. 0.35, which is lower than the proportion of unbalanced trees ($\frac{2}{3}$) expected under frequent sex.

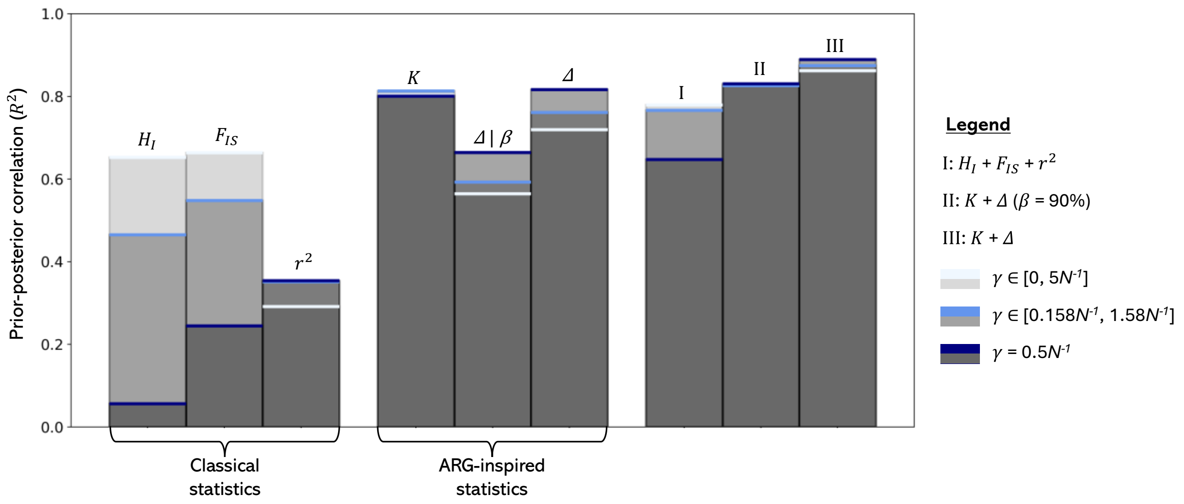


**Fig. S12.** Genealogical statistics outperform classical statistics in predicting the rate of sex across the simulated scenarios. As a measure of how well each statistic, or summary vectors composed of multiple statistics (combinations **I-III**), predicted $\sigma$, we used the coefficient of determination ($R^2$) between the prior and posterior values obtained through rejection-based approximate Bayesian computation (R package 'abc', tolerance of 0.01, log-transformed parameter values) (Csilléry et al., 2012). For each simulated scenario, we designated 10 replicates as 'observed' and the remaining 90 as draws from the prior distribution. We repeated the procedure for progressively restricted ranges of $\gamma$ values (see legend). Overall, the highest $R^2 = 0.89$ was achieved by a combination of $K$ and $\Delta$ (no bias; $\gamma = 0.5N^{-1}$), whereas among classical statistics the highest $R^2 = 0.78$ was achieved by a combination of $H_I$, $F_{IS}$ and $r^2$ ($\gamma \in [0, 5N^{-1}]$).

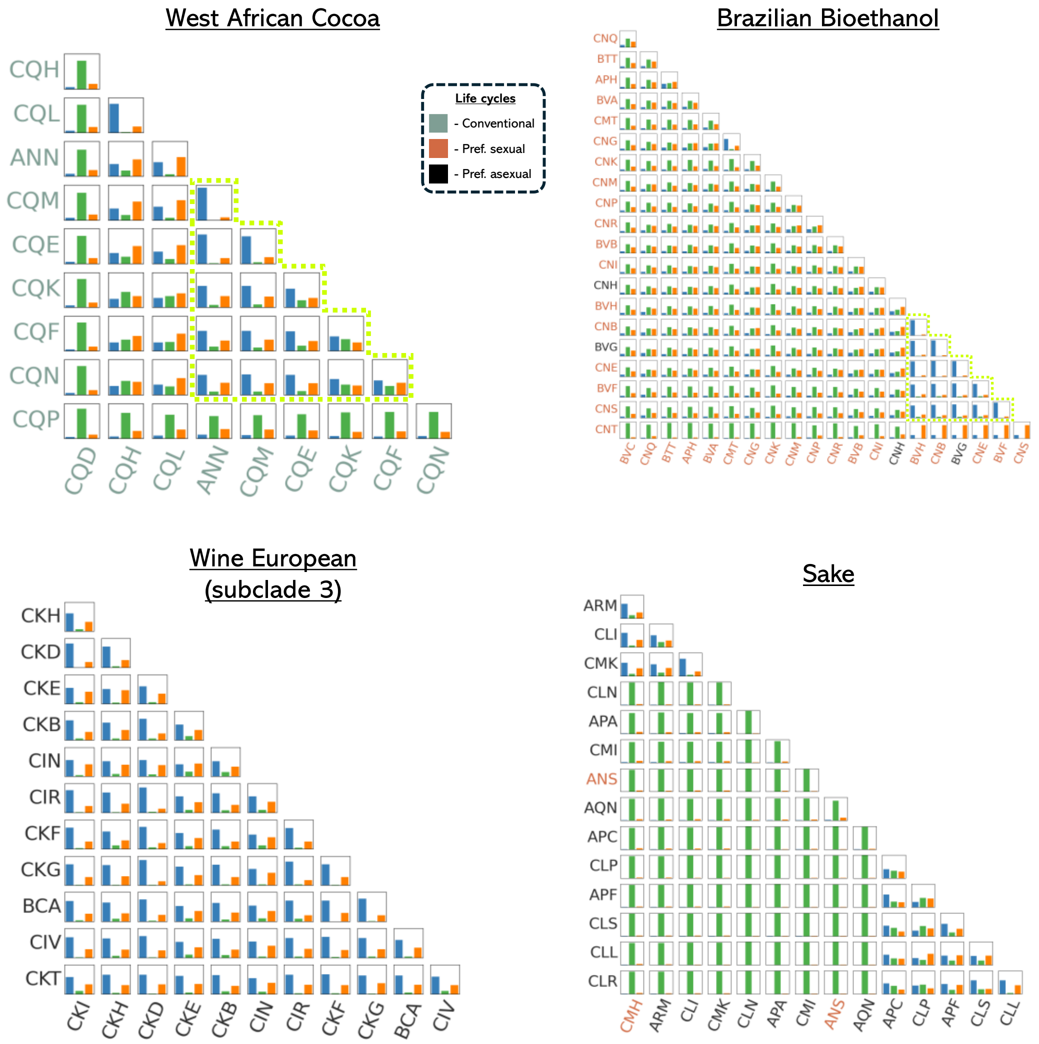


**Fig. S13.** Tree composition per pair of isolates in the studied *S. cerevisiae* clades. Isolate codes are coloured according to life cycle (Becerra-Rodríguez et al., 2026). Yellow outlines demark putative asexual sub-clades with 0-skewed *Δ* values (see Fig. 5B and Fig. S13). While Fig. 5B shows a subset of Brazilian Bioethanol isolates for clarity, all 21 isolates are included here.

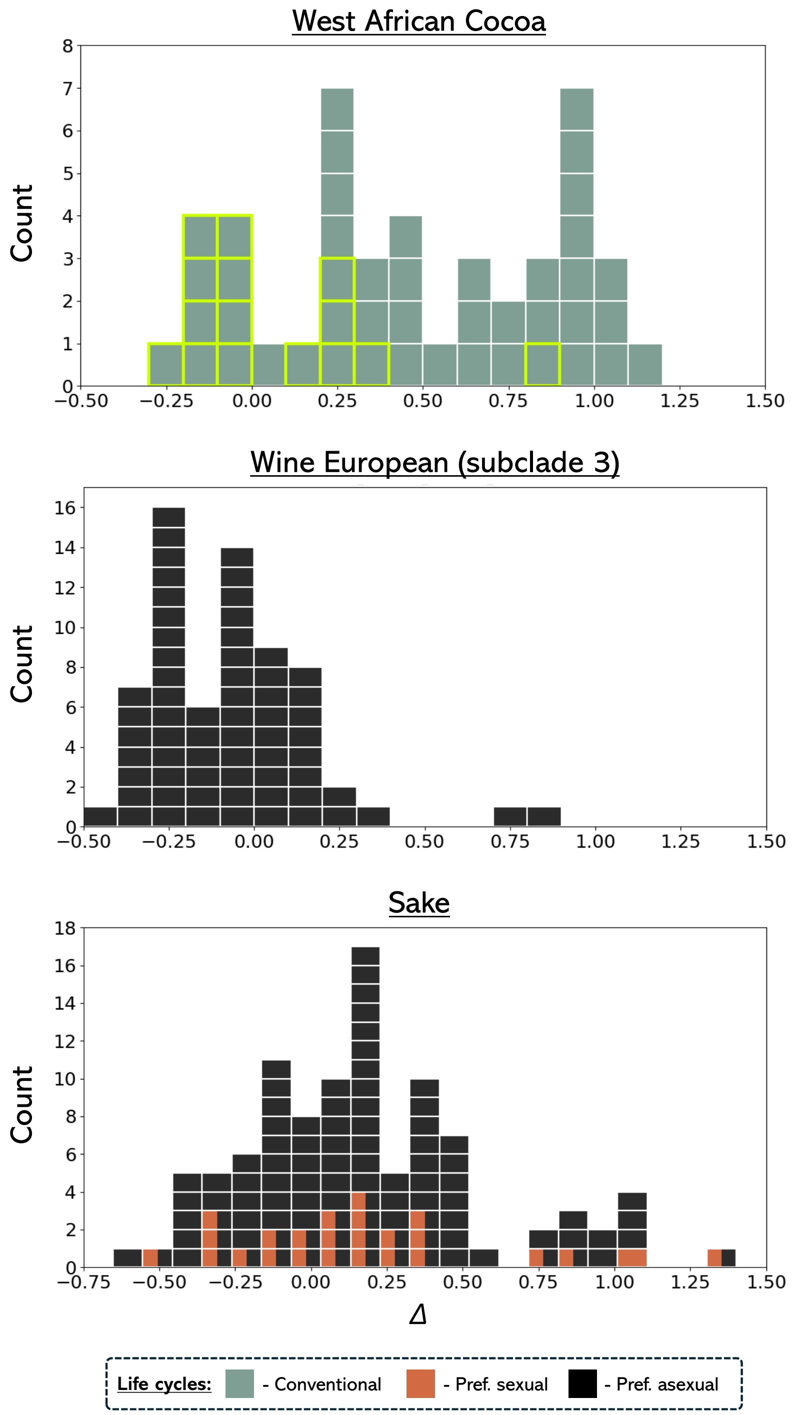


**Fig. S14.** Distribution of *Δ* values among pairs of isolates in the studied *S. cerevisiae* clades (see Fig. 5B for Brazilian Bioethanol). In the histograms, each *Δ* value is represented by a block coloured according to life cycle assignments of the two isolates (left vs right). Yellow outline demarks a putative asexual in subclade in the West African Cocoa clade (compare with Fig. S13).

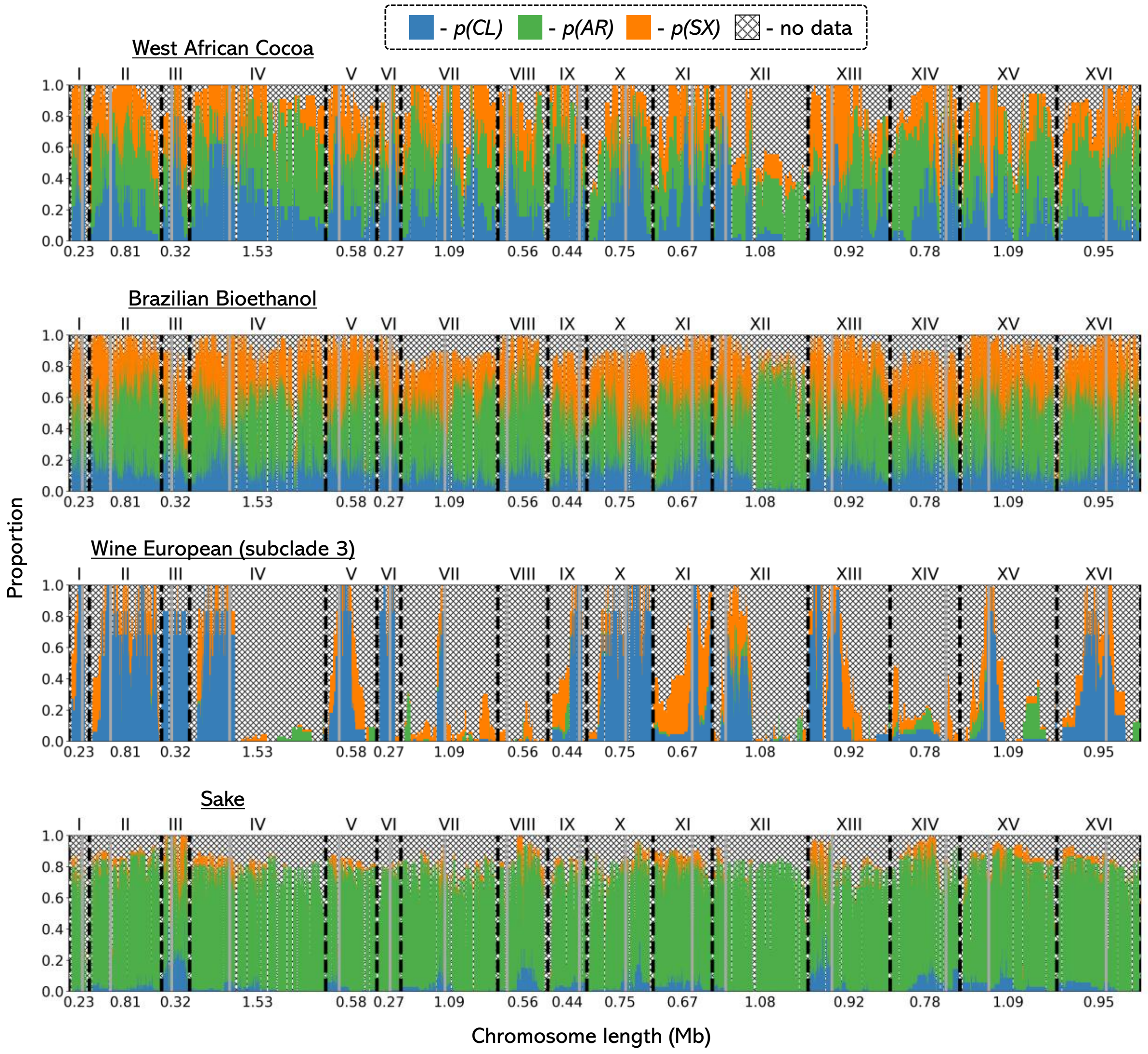


**Fig. S15.** Tree composition across the 16 chromosomes in the studied *S. cerevisiae* clades. Centromeres are marked with grey lines; chromosome boundaries are marked with dashed, black lines. The ‘no data’ category includes multiple sources of missingness, including alignment gaps, polytomies (excluded from tree composition analysis), and regions lacking informative SNPs.

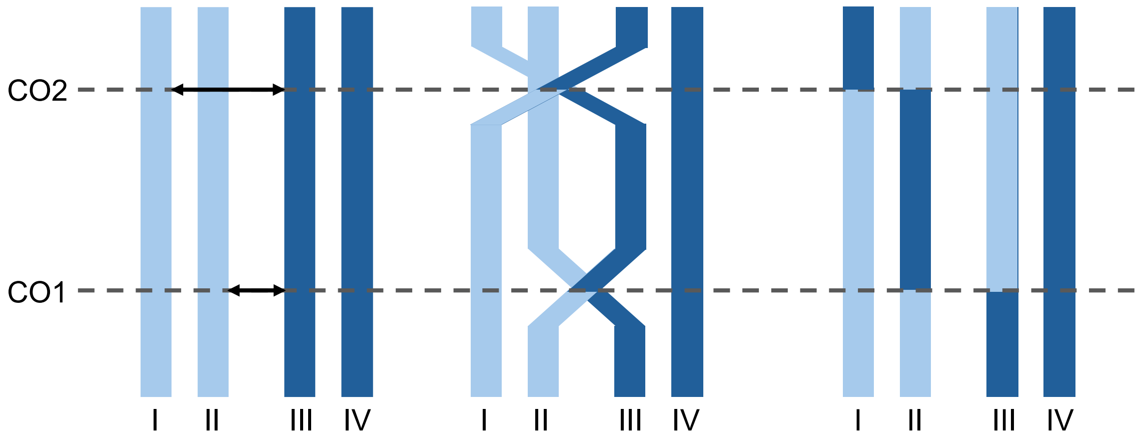


**Fig. S16.** Resolution of crossovers. Two crossovers occur: between chromatids II and III (CO1) and between chromatids I and III (CO2). Although chromatid II was not involved in CO2, the reconnecting of broken DNA molecules results in a double crossover on this chromatid.

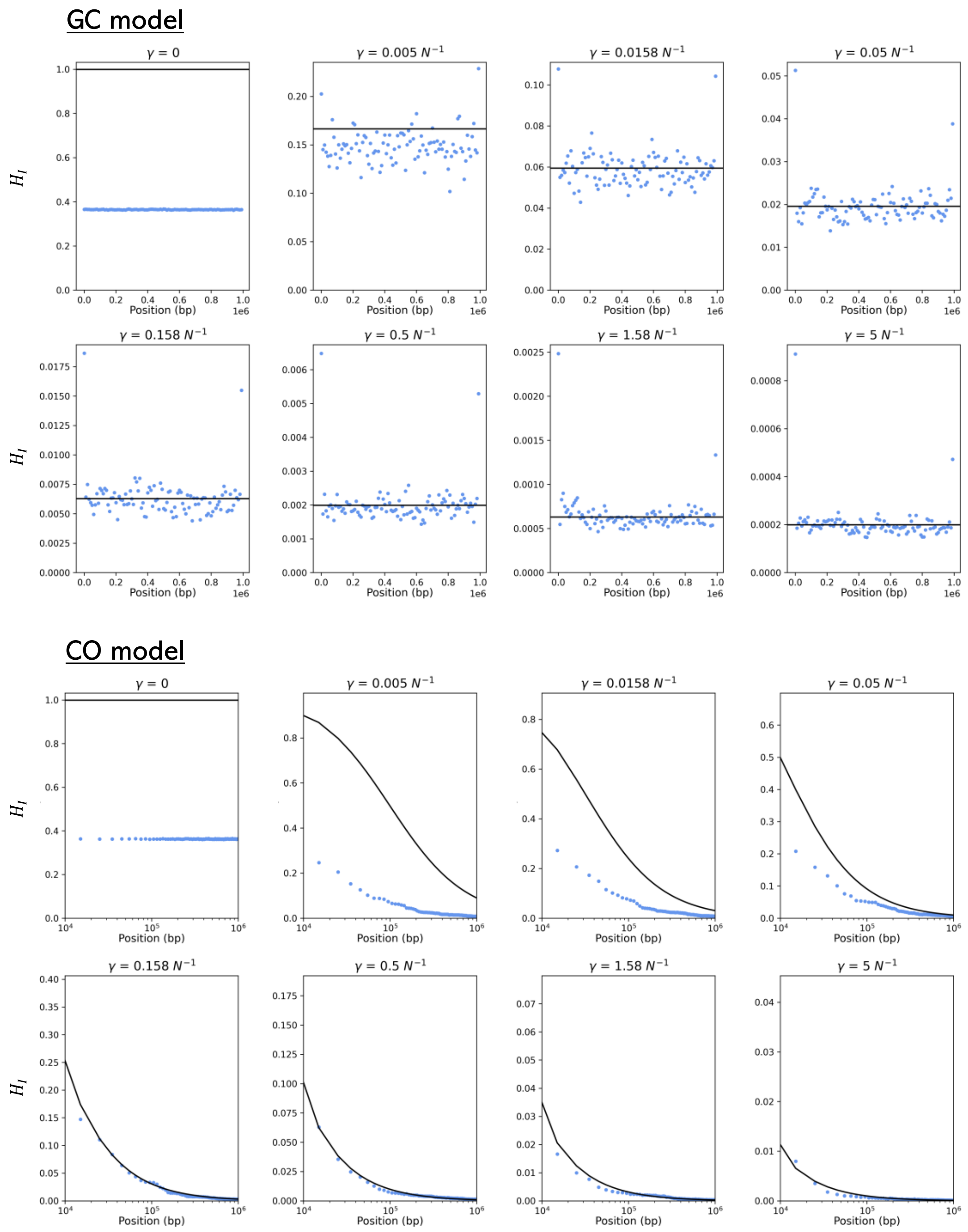


**Fig. S17.** $H_I$ after $5 \times 10^5$ simulated generations of obligate asexuality (dots) compared against the analytical predictions (lines; see section A, Eq. 2 and Eq. 7 for the GC and CO expressions, respectively).

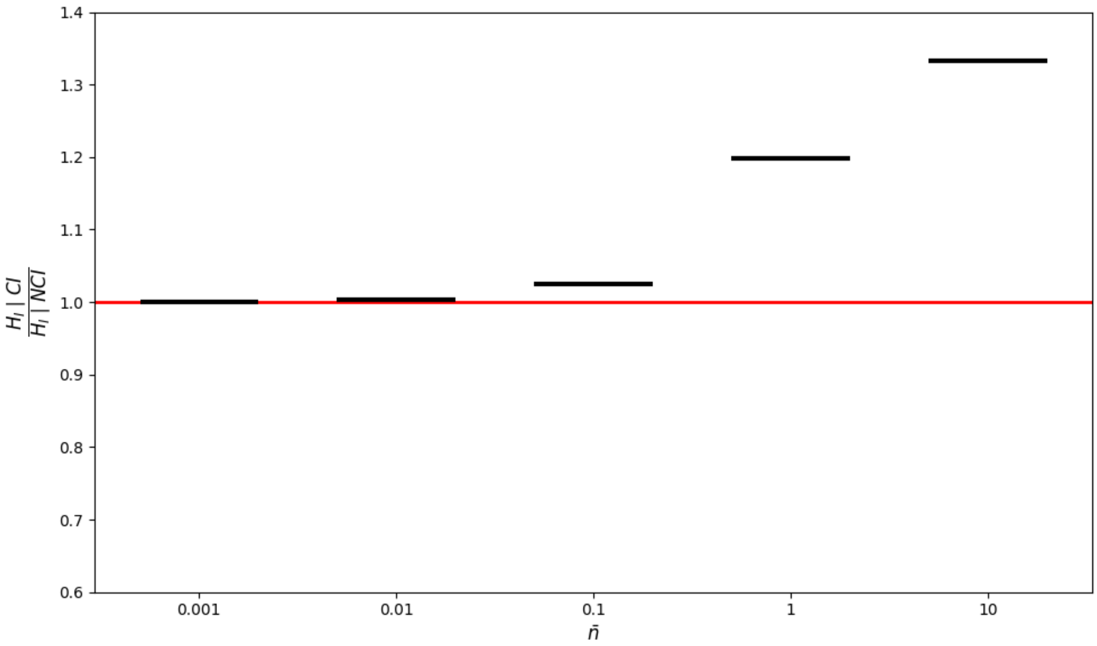


**Fig. S18.** Comparison of the CI and NCI models of crossover recombination under the CO model of asexuality. A ratio of the equilibrium individual-level heterozygosity $\widehat{H}_I$ under the CI model to that expected under the NCI model as a function of the expected number of crossovers between the centromere and the focal locus $\bar{n}$.

### Supporting Information Tables

**Table S1.** Simulation parameters and their values used in the simulation pipeline.

| Parameter | Definition | Values |
|---|---|---|
| $L$ | Chromosome length in bp | $10^6$ |
| $N$ | Diploid population size | $10^3$ |
| $\sigma$ | Probability of offspring produced sexually by hermaphroditic parents | [0, $10^{-5}$, $3.162 \times 10^{-5}$, $10^{-4}$, $3.162 \times 10^{-4}$, $10^{-3}$, $3.162 \times 10^{-3}$, 0.01, 0.03162, 0.1, 0.3162] |
| $r_{GC}$ | Per-site per-generation rate of gene conversion tract initiation under the GC model | [0, $10^{-9}$, $3.162 \times 10^{-9}$, $10^{-8}$, $3.162 \times 10^{-8}$, $10^{-7}$, $3.162 \times 10^{-7}$, $10^{-6}$] |
| $\lambda$ | Mean gene conversion tract length in bp; tract length is geometrically distributed where $\frac{1}{\lambda}$ is the probability of tract termination | $5 \times 10^3$ |
| $r_{CO}$ | Per-site per-generation rate of crossing-over under the CO model | [0, $2.019 \times 10^{-11}$, $6.324 \times 10^{-11}$, $2 \times 10^{-10}$, $6.326 \times 10^{-10}$, $2.002 \times 10^{-9}$, $6.345 \times 10^{-9}$, $2.02 \times 10^{-8}$] |
| $\gamma$ | Mean per-site per-generation rate of loss of heterozygosity (LOH) | Section A: Eq. 2 (GC model) and Eq. 7 (CO model) |
| $r$ | Per-site per-generation rate of crossing-over during sexual reproduction | $10^{-6}$ |
| $\mu$ | Per-site per-generation mutation rate | $5 \times 10^{-7}$ |

**Table S2.** Comparison of genealogical reconstruction methods.

| Criterion | sticcs (Martin, 2026) | SINGER (Deng et al., 2025) |
|---|---|---|
| **General framework** | Heuristic, based on parsimony. | Bayesian, model-based ARG inference using MCMC sampling. |
| **Assumptions** | Minimal assumptions – infinite-sites mutation model (Kimura, 1969) and perfect phylogeny. Not based on an explicit demographic model. | Sequentially Markov coalescent (McVean & Cardin, 2005); parametrised by input mutation/recombination rates. |
| **Local tree inference** | Local tree topologies reconstructed deterministically from collections of mutually compatible sites – all trees supported by mutational information. | Local tree topologies inferred probabilistically using a hidden Markov model and posterior sampling – trees often without local mutational support. |
| **Coalescence time inference** | No inference of coalescence times. | Coalescence times inferred probabilistically using a hidden Markov model and posterior sampling. |
| **Recombination breakpoint inference** | Breakpoints inferred from incompatibilities between sites (four-gamete test logic; Hudson & Kaplan, 1985). | Recombination events inferred probabilistically within the Bayesian framework. |
| **Output** | A single inferred ARG (deterministic reconstruction). | Posterior samples of ARGs. |

**Table S3.** Estimates of the rate of LOH per site ($\gamma$) from mutation accumulation (MA) experiments. We report the total estimates of $\gamma$ which include contributions from short-range (e.g., gene conversion) and long-range (e.g., crossing-over, break-induced replication) forms of asyngamous recombination.

| Organism | Reference | Detection method | Mechanism of asexuality | Estimate of $\gamma$ (LOH $site^{-1}$) |
|---|---|---|---|---|
| *Daphnia pulex* | Omilian et al. (2006) | Microsatellite analysis | meiotic | $2.1 \times 10^{-4}$ |
| | Xu et al. (2011) | | | $3.9 \times 10^{-5}$ |
| | Keith et al. (2016) | WGS | | $1.18 \times 10^{-5}$ |
| | Flynn et al. (2017) | | | $4.8 \times 10^{-5}$ |
| *Adineta vaga* | Houtain, Derzelle, Lliros, et al. (2024) | WGS | meiotic | $3.24 \times 10^{-4}$ |
| *Saccharomyces cerevisiae* | Sui et al. (2020) | WGS | mitotic | $2.6 \times 10^{-5}$ |
| | Dutta et al. (2021) | | | $7.1 \times 10^{-5}$ |

**Table S4.** The studied clades in the context of the *Saccharomyces cerevisiae* genomic dataset of Peter et al. (2018) and Loegler et al. (2025). Using short-read data generated by Peter et al. (2018), and additional long-read data, Loegler et al. (2025) generated chromosome-level, phased assemblies for selected isolates. For the present analysis, only diploid, euploid and heterozygous isolates (as classified by the two studies) were considered 'targets'; however, phased assemblies were not generated for all such isolates. A heterozygosity bias (*sensu* Main Text) remains an important consideration when interpreting genealogical signatures recovered from the subset of genomes that were phased.

| Clade | Peter et al. (2018) | | | | | Phased in Loegler et al. (2025) |
|---|---|---|---|---|---|---|
| | Total | Diploid | Euploid | Het.* | Target | |
| West African Cocoa | 13 | 13 | 13 | 13 | 13 | 10 |
| Brazilian Bioethanol | 35 | 35 | 32 | 26 | 24 | 21 |
| Wine European (subclade 3) | 24 | 24 | 21 | 24 | 21 | 12 |
| Sake | 47 | 47 | 20 | 38 | 14 | 15** |

*Heterozygous.

**The CLN isolate was classified as homozygous by Peter et al. (2018), but as heterozygous in Loegler et al. (2025).
